# CellVoyager: AI CompBio Agent Generates New Insights by Autonomously Analyzing Biological Data

**DOI:** 10.1101/2025.06.03.657517

**Authors:** Samuel Alber, Bowen Chen, Eric Sun, Alina Isakova, Aaron J. Wilk, James Zou

## Abstract

Modern biology increasingly relies on complex, high dimensional datasets such as single-cell RNA sequencing (scRNA-seq). However, the richness of such data means that conventional analyses may only scratch its surface. Extracting meaningful insights from these datasets often requires advanced computational methods and domain expertise. Current AI agents for biology are primarily focused on executing user-specified commands and are therefore limited by the user’s creativity and familiarity with which kinds of analyses are useful. Furthermore, these agents do not account for prior analyses already attempted by researchers, reducing their ability to build upon existing work. To address these limitations, we introduce CellVoyager, an AI agent that autonomously explores scRNA-seq datasets in novel directions conditioned on prior user-ran analyses. Built on large language models, CellVoyager ingests both the dataset and a record of prior analyses to generate and test new hypotheses within a Jupyter notebook environment. We evaluate CellVoyager on CellBench, a new benchmark based on 50 published scRNA-seq studies encompassing 483 analyses. Given only the background sections of these papers, CellVoyager outperformed GPT-4o and o3-mini by up to 20% in predicting which analyses the authors eventually conducted. We then carried out three in-depth case studies where CellVoyager is given previously published papers with their scRNA-seq datasets and conducts analyses to generate new findings. The original authors of each study evaluated these findings and consistently rated them as creative and sound; 80% of the agent’s hypotheses were deemed scientifically interesting. For example, in one case study, the agent found that CD8^+^ T cells in COVID-19 infection are more primed for pyroptosis, which was not explored by the original researchers. CellVoyager also reanalyzed a brain aging dataset to discover a previously unreported association between increased transcriptional noise and aging in the subventricular zone of the brain. These results demonstrate that CellVoyager can act both autonomously and collaboratively to accelerate hypothesis generation and computational biology. It also highlights the potential of agents like CellVoyager to unlock new biological insights by reanalyzing the vast existing biological data at scale.

## 1 Introduction

Modern biological research often involves collecting and analyzing complex, high-dimensional datasets. These datasets range from multi-omics assays to spatial and temporal profiling, each harboring rich biological insights [2, 25, 37]. Yet, the high dimensionality that enriches these datasets also introduces substantial analytical challenges, requiring complex, domain-specific computational methods in order to extract real biological signal [7, 37, 47, 49]. Many of these methods have steep learning curves, especially for researchers without computational backgrounds. Furthermore, there are numerous biological hypotheses that can formalized using the various combinations of features in these high-dimensional datasets. Many of these hypotheses may remain unexplored because of the vast scale of this space of analyses and the expertise necessary to execute some of these analyses.

Recently, large language models (LLMs) have demonstrated their knowledge across various biological domains [22, 42] and for writing complex code [17]. In particular, there is growing interest in LLM-based AI agents, with prior works demonstrating the ability of these agents to take natural language instructions from users and automatically generate, execute, and explain relevant code [51, 56, 59]. These capabilities can potentially lower the barrier to entry for many non-computational researchers who want to leverage computationally intensive data modalities. Beyond following instructions, AI agents have demonstrated potential in helping researchers navigate the complex landscape of possible analyses by autonomously generating hypotheses [15, 18, 28, 32] and applying diverse analytical methods [34, 46, 52] to uncover insights.

In a typical biological research setting, investigators have a set of analyses they plan to try or have already executed. Therefore, a truly collaborative AI agent should be able to take in this set of analyses and either build upon them or propose new directions that extend upon the prior work, complementing the investigator’s prior efforts rather than duplicating them. Although recent LLM-based agents have shown promise in scientific hypothesis generation [12, 15, 18, 28, 32], it remains untested if they can reliably generate analyses meaningfully distinct from an inputted set of analysis ideas.

Single-cell RNA sequencing (scRNA-seq) provides an ideal testbed for these challenges. By measuring gene expression at a single-cell resolution, this type of sequencing has been used to identify pathogenic cell subpopulations, infer cell-state transitions, and uncover differentially expressed genes across a condition of interest [5, 8, 9]. Due to the high dimensionality of scRNA-seq data and the presence of various technical and biological confounders, analyzing it often requires substantial computational expertise [24]. This has led to the development of thousands of open-source computational tools for analyzing scRNA-seq data [57]. Moreover, the extensive gene coverage and high resolution of scRNA-seq data enables the formulation and exploration of a large set of biological hypotheses. Collectively, these result in a vast space of possible computational analyses that can be conducted on the data. Depending on the time, resources, and expertise available to a set of researchers, they may be limited to a narrow subset of these analytical workflows, potentially missing important biological findings. AI agents capable of autonomous dataset exploration could help address these issues by leveraging the extensive biological and computational knowledge of their underlying LLMs to explore a wide range of analyses. This autonomous exploration could also help find biological insights that are counterintuitive and that a researcher may not think to test. While recent work has begun to explore AI-generated hypotheses in general experimental settings [39] or execution of single-cell analyses given user commands [52, 55, 58, 60], fully autonomous dataset exploration remains largely unaddressed in the context of scRNA-seq data or other high-dimensional biological datasets.

In this work, we introduce CellVoyager: an LLM-driven AI agent that autonomously explores scRNA-seq datasets. By taking in the processed scRNA-seq dataset, biological background information, and a record of analyses already performed on the dataset, CellVoyager generates and executes new analytical pipelines that build directly on prior work. Unlike earlier systems, we focus on how well CellVoyager can complement and extend an existing set of analyses, facilitating a more collaborative human-AI research workflow and enabling researchers to explore analyses beyond their areas of expertise. This work focuses on using published research papers as input into CellVoyager, since this represents a challenging setting where extensive effort has been put into exploring the dataset. In this context, the paper contains both biological background necessary to help inform which analyses the user cares about and the past analyses the agent should build upon or avoid. To demonstrate the utility of CellVoyager, we applied it to a collection of published scRNA-seq studies, where some of the original authors of the respective manuscripts evaluated the agent’s findings and suggestions.

## 2 Related Work

### Applications of LLMs in Bioinformatics

Large language models (LLMs) have increasingly demonstrated their potential across a broad range of domains within bioinformatics. One popular research direction involves applying LLM architectures to biological sequences. These efforts include modeling and generating insights from DNA [6, 35], protein [26], and RNA sequences [10], treating these sequences as a “language” of biology. Beyond sequence modeling, another promising area is extracting biological knowledge from a pretrained or fine-tuned LLM. This could include directly querying an LLM fine-tuned on biomedical literature [16] or something more applied, like extracting biologically meaningful embeddings from free-text gene descriptions [11]. Such methods provide useful analytical tools for domain-specific downstream applications.

### LLM Agents for Bioinformatics

An emerging paradigm in using LLMs for bioinformatics is the creation of autonomous or semi-autonomous LLM-based agents for performing complex scientific workflows. Rather than relying on a specialized tool or model to solve a specific task, these agents typically integrate multiple domain-specific tools and interface with natural language commands, being able to analyze biological data based on human prompt or guidance. Notable examples include agents designed for analyzing gene expression datasets [27], drug discovery [3], pathology images [30], spatial transcriptomics [46], and gene perturbation experiments [39]. These systems demonstrate the ability of LLMs to chain together specialized tools to analyze bioinformatics data.

### LLM Agents for Single-Cell Transcriptomics

A particularly impactful subfield for LLM agents in bioinformatics is the application to single-cell RNA sequencing analysis. Datasets in this domain are often high-dimensional and require complex processing pipelines as well as the use of domain-specific tools. Recent works such as CellAgent [52], CompBioAgent [58], AutoBA [60], and BIA [55] demonstrate the feasibility of LLM agents to execute user commands for scRNA-seq data analysis tasks. These agents are primarily designed to automate the data analysis process, reducing the need for a bioinformatician to manually analyze data using domain-specific tools and packages. Using these agents, a user without technical background can input intuitive natural language instructions and rely on the agent to conduct the analysis. While these agents have demonstrated the ability to carry out some standard scRNA-seq tasks, such as batch correction [52] and cell type annotation [52, 58], they do not formulate nor explore their own hypotheses. Therefore, they are limited to executing specific user requests. In contrast, CellVoyager is capable of autonomously exploring the single-cell dataset and propose, as well as conduct, its own analyses. Additionally, CellVoyager can be conditioned on analyses that the user has attempted previously and conduct further analyses that can either complement what the user has tried or call attention to underexplored aspects of the data. This feature is especially valuable in the context of single-cell transcriptomics, where important biological signals may be overlooked when manually analyzing such complex and high-dimensional datasets.

## 3 CellVoyager Methodology

CellVoyager is an agent-based framework that autonomously explores scRNA-seq datasets while incorporating the user’s past analyses to prevent redundancy and help find analyses the user may be unfamiliar with. This approach enables researchers to uncover biologically meaningful insights that may lie outside their immediate expertise while simultaneously evaluating the creativity and problem-solving capacity of LLMs in a data-driven research setting. By integrating an LLM with live code execution within a Jupyter environment, CellVoyager dynamically generates, iteratively refines, and executes novel analysis plans—termed “exploration blueprints.” See **Figure 1** for a schematic of the CellVoyager agentic workflow and **Section A.1** for a more detailed description of the agent, including the prompts used.

**Figure 1:**
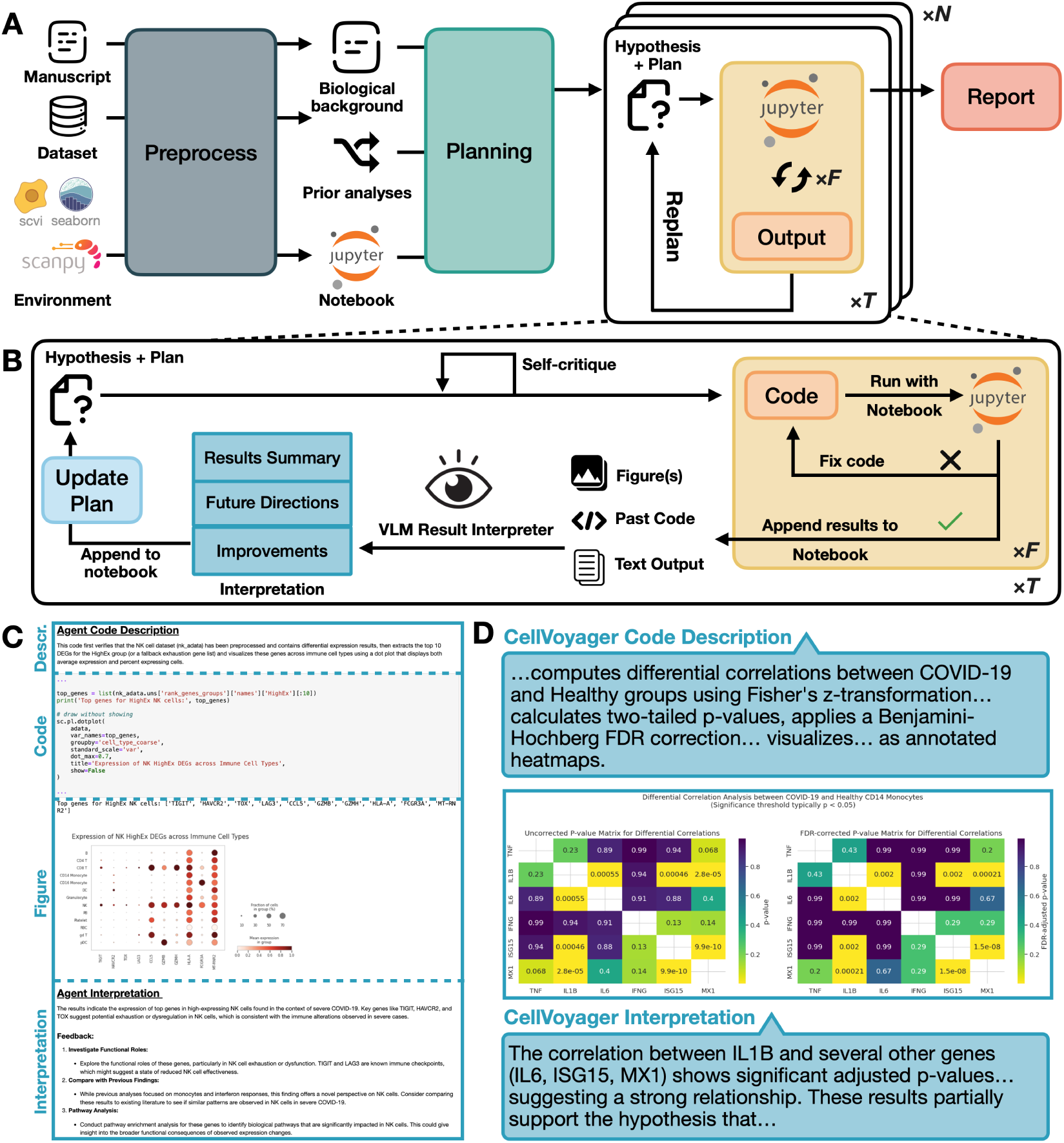
Schematic of the CellVoyager agentic framework. **A. Overall schematic of CellVoyager** (1) The manuscript text, associated scRNA-seq data, and a Python environment specification are given as input. The biological background and prior analyses performed in the manuscript are extracted and a Jupyter notebook is initialized. (2) Exploration blueprints (consisting of a hypothesis and a stepwise plan) are generated. The agent self-critiques this blueprint, incorporating any suggested improvements. (3) Each hypothesis is explored for T steps. Each step involves generating and executing code, with up to F steps of bug fixing, replanning based on the code execution output, and self-critiquing the new plan. This process can be repeated for N distinct hypotheses, with the agent being given past hypotheses it tested to prevent it from repeating the same analyses. A final report is generated that summarizes noteworthy findings from the N exploration trajectories. **B. Overview of CellVoyager analysis and interpretation module.** The self-critiqued plan contains python code for the next step of the analysis. This code is executed and the outputs from running the code, along with the contents of past code cells, are fed into a vision-language model (VLM) for interpretation. The VLM outputs a summary of the outputs, suggested future directions based on promising results, and possible ways to improve the analysis; these inform how the agent decides to choose the next step in the analysis and CellVoyager updates its analysis plan accordingly. **C. Basic analysis block of CellVoyager** in a Jupyter notebook, consisting of a description of the analysis step, the code to carry out the analysis, the outputs and figures, and the interpretation of the outputs and figures. **D. An example of an analysis block** of CellVoyager, with code omitted for brevity.

### 3.1 Agent Setup

CellVoyager requires two primary inputs: a processed scRNA-seq dataset and a report that explains the biological context of the dataset (and previous analyses, if there are any). The report provides context to the biological problem, details what is in the single-cell dataset, and describes the prior computational single-cell analyses performed on the dataset. In this work, we use a published manuscript as the report because of the challenge in finding unexplored analytical directions in them. To guarantee reproducibility and prevent dependency conflicts, CellVoyager operates within a fixed Jupyter kernel that includes popular single-cell packages part of the scverse [45] like scanpy and scVI [29, 50]. These packages’ names are also inputted into the agent to inform it which packages it has access to in its environment.

The agent first generates a summary of the paper, separating it into (1) biological background, (2) analyses attempted, and (3) details about the dataset used. Moving forward, CellVoyager uses this summary rather than the full text of the paper. The agent also generates a Jupyter notebook, initializing the execution environment by loading in the dataset and the relevant python packages. By interfacing with an interactive notebook, CellVoyager is able to access previous results, data, and intermediate outputs, allowing it to efficiently iterate on its analysis.

### 3.2 Analysis planning, Execution, and Replanning

The central purpose of CellVoyager is to find a single-cell analysis distinct from the ones conducted in the paper but still biologically interesting and in line with the research goals of the paper. The agent operates through creating and reflecting upon “exploration blueprints”. These blueprints have two main components: a hypothesis and a step-by-step analysis plan. This blueprint provides the high-level plan for the agent to execute the analysis.

Given a blueprint, the agent iterates on its analysis plan with a budget of T iterations. At each iteration, CellVoyager reflects on its current exploration blueprint (by self-critiquing it), modifying the blueprint as necessary, and executes its analysis in the form of Python code appended as a code cell to a Jupyter notebook. If the code produces an error, the agent rewrites the code for up to F fix attempts. If the code runs successfully, CellVoyager interprets the images and text outputted by the code via a vision language model. From its interpretation of these results, the agent then replans the next steps in its current analysis as needed. For each execution output, the agent also provides a natural language interpretation of the results and appends it as a markdown block to the notebook. If the code errors cannot be fixed within the allowed attempts, then the agent replans the next steps based on the error messages, formulating alternative analytical trajectories. See **Figure 1A,B** for the process visualized.

In the Jupyter notebook, at each step, CellVoyager’s basic analysis block consists of a description of its current action, a code cell executing that action, the outputs from running the code cell, and an interpretation of the outputs. An example of a CellVoyager-generated analysis is shown in **Figure 1C,D**, which includes the agent’s interpretation of the figure.

CellVoyager runs analyses sequentially, where each analysis is conditioned on the previous analyses to ensure CellVoyager doesn’t repeat analyses. After all analyses are complete, the agent extracts the most promising results and writes a report detailing its hypotheses, procedure, and interpretation of those results. This allows the agent to effectively explore many analyses and then highlight a subset that are scientifically interesting to the user. Since the agent’s full exploration trajectory is documented in the Jupyter notebook, with headers detailing the agent’s hypothesis and results interpretation at every iteration, the notebook acts as a detailed and interpretable full report if the user desires more detail.

## 4 Evaluating CellVoyager Analysis Generation with CellBench

To evaluate the ability of CellVoyager to propose relevant analyses for a research topic, we propose CellBench: a benchmark for measuring the ability of an agent to infer missing analyses in a single-cell transcriptomics research paper. Each query in CellBench is based on a published study that analyzes single-cell transcriptomic data. For each study, we use an off-the-self LLM (gemini-2.5-pro-preview) to extract the biological background from the paper as well as a list of all of the single-cell analyses reported in the paper. The biological background section acts as the input context for each CellBench query. Given this context, the agent is tasked to propose single-cell analyses that it thinks the paper should perform. The analyses actually performed in this paper act as the ground truth analyses. The overlap between the proposed analyses and the ground truth analyses are measured and used as the evaluation metric. To ensure the benchmark is scalable, we use an LLM judge for determining whether a given analysis shows up in a set of ground truth analyses. See **Figure 2A** for a schematic of the curation process. To get a sense of how accurate the LLM judge is, two PhD students independently evaluated o3-mini’s outputs on a subset of 47 generated analysis ideas; the concordances rates between each rater and the LLM judge were 89% and 85% respectively, indicating that the LLM judge is a reliable heuristic.

**Figure 2:**
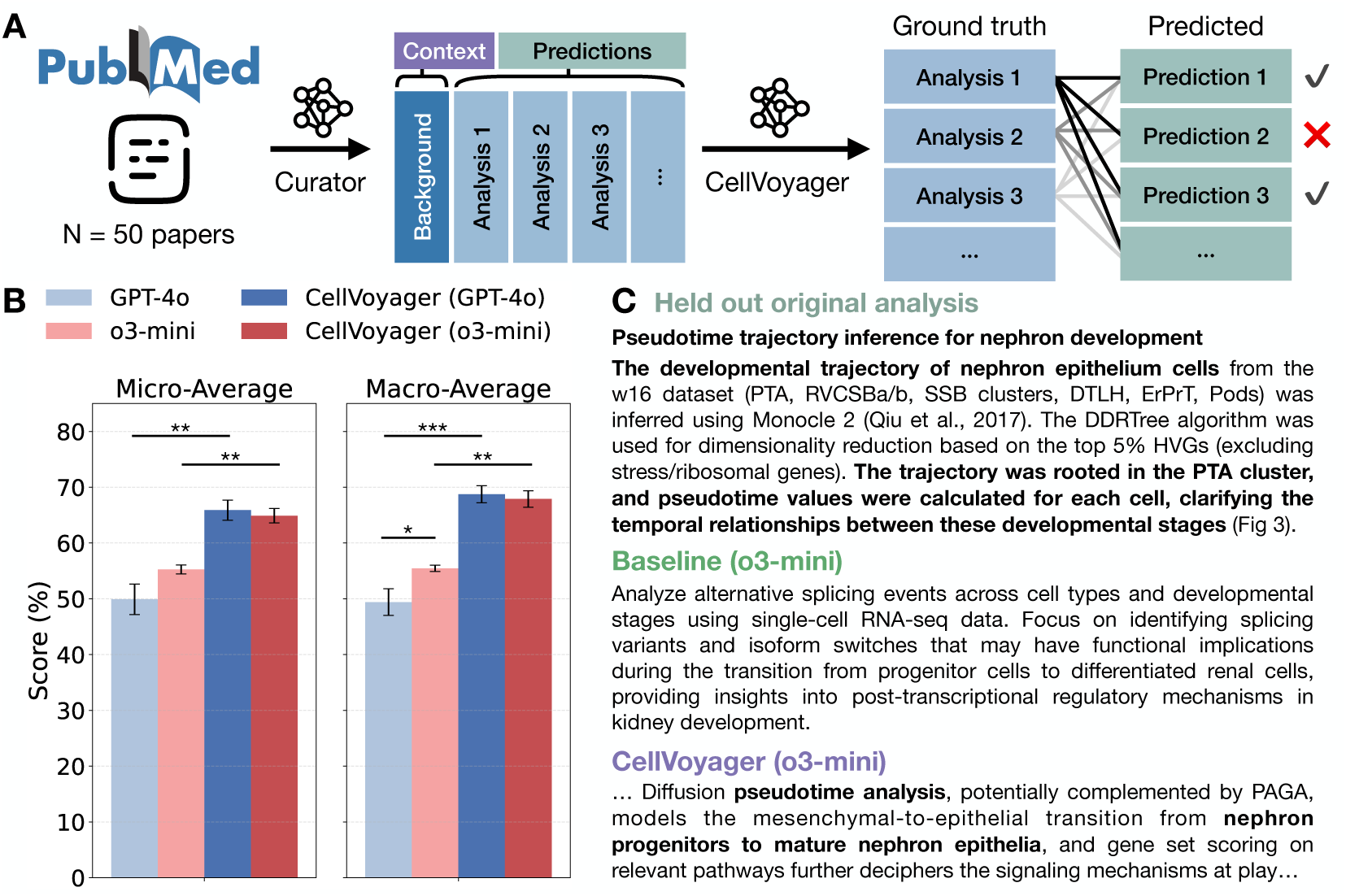
CellBench evaluation. **A. Benchmark Curation.** CellBench is curated from 50 papers that focus on analyzing scRNA-seq data. An LLM extracts the biological background information and enumerates the independent analyses performed in the paper. The agent is tasked with predicting analyses only given the paper’s background section and is evaluated on how accurately it predicts the paper’s held-out analyses. **B. Performance of base models and CellVoyager**. With both LLMs, CellVoyager outperforms the base LLM model and CellVoyager (GPT-4o) had the highest accuracy. Bar values represent the average accuracy score in predicting a correct held-out analysis. Error bars represent standard deviation from running the benchmark three times. ∗: *p* < 0.05,^∗∗^: *p* < 0.01,^∗∗∗^: *p* < 0.001. **C. Example CellBench query and agent response.** In this example, o3-mini (Baseline) proposes an analyses irrelevant to the paper. CellVoyager (using o3-mini as its LLM) proposes an analyses that matches one of the analyses performed in the paper more closely (see bolded text).

Overall, CellBench is derived from 50 published works, consisting of 483 total analyses. Given the background section of a paper, a LLM or agent is tasked with predicting the analyses conducted by the authors of the paper. Since CellVoyager includes additional modules that write and correct code, for a fairer comparison with baselines, we only run the initial idea generation phase of CellVoyager. There, the agent finds an analysis plan and reflects upon it to find potential improvements. Additionally, we adjusted the prompt in CellVoyager to propose the most probable held-out analyses rather than solely focusing on outputting unique analyses since the latter is biased towards more unconventional analyses. CellBench does not task the agent to write code to carry out the proposed analyses.

We evaluated CellVoyager using different LLMs as well as the base LLMs by themselves on CellBench. For all configurations, we use the default parameters from the OpenAI API endpoint and reasoning effort of medium for o3-mini. We report both the micro-averaged and macro-averaged accuracies across the analyses pertaining to each paper. See **Figure 2B,C** for results and an illustrative example. We find that CellVoyager outperforms the base LLM by a considerable margin. For the best performing case, CellVoyager using GPT-4o as the engine, the agent outperforms GPT-4o by 16% in terms of micro-averaged accuracy (p < 0.01) and 19.33% in terms of macro-averaged accuracy (p < 0.001). Although CellVoyager using o3-mini underperforms compared to using GPT-4o, the difference is not statistically significant (p ≫ 0.05). Since o3-mini has been shown to be better in the latest coding benchmark leaderboards [20, 48], we base our agent on o3-mini in our case studies in the next section.

## 5 Computational Biology Case Studies

To evaluate whether CellVoyager can meaningfully extend high-quality research, we tested its ability to build upon peer-reviewed and published single-cell transcriptomics research papers. These are scenarios in which considerable effort has already been devoted towards analyzing a single-cell dataset. Therefore, it should be difficult for an agent to find interesting results related to the paper’s core goals that were not already mentioned in the paper. A complicating factor is that some analyses may have been conducted but were omitted from the final manuscript. To control for this, we included one of the authors of each paper in the evaluation process, ensuring familiarity with both the published analyses and any potential internal results not included in the paper.

We select three published single-cell transcriptomic studies, spanning several diseases and tissue types [7, 47, 49]. For each case study, CellVoyager generates eight distinct analyses, each encapsulated in a standalone Jupyter notebook. Each analysis’ code and CellVoyager’s reasoning trace is documented within a standalone Jupyter notebook, typically containing 5-8 analysis steps. We selected the five analyses with the highest percentage of successfully executed code cells for further evaluation (randomly choosing among ties). We provided these Jupyter notebooks to an author of each respective paper along with an evaluation rubric (**Appendix Figure 1**). For each analysis, reviewers were asked to assess CellVoyager’s creativity (on a scale from one to four; this measures how distinct the analysis is from those done in the paper), whether the analysis was biologically useful, whether the correct methods were used, and if an interesting hypothesis was raised by the agent that the author would like to explore further. To assess CellVoyager’s capacity for human-guided iteration, we include the reviewer’s feedback as input into CellVoyager for a subset of analyses. The agent then generated revised versions of these analyses via incorporating the reviewers’ suggestions for how to improve the analyses. A summary of the evaluation results are shown in **Figure 3** with the proceeding sections going into more detail on each case study.

**Figure 3:**
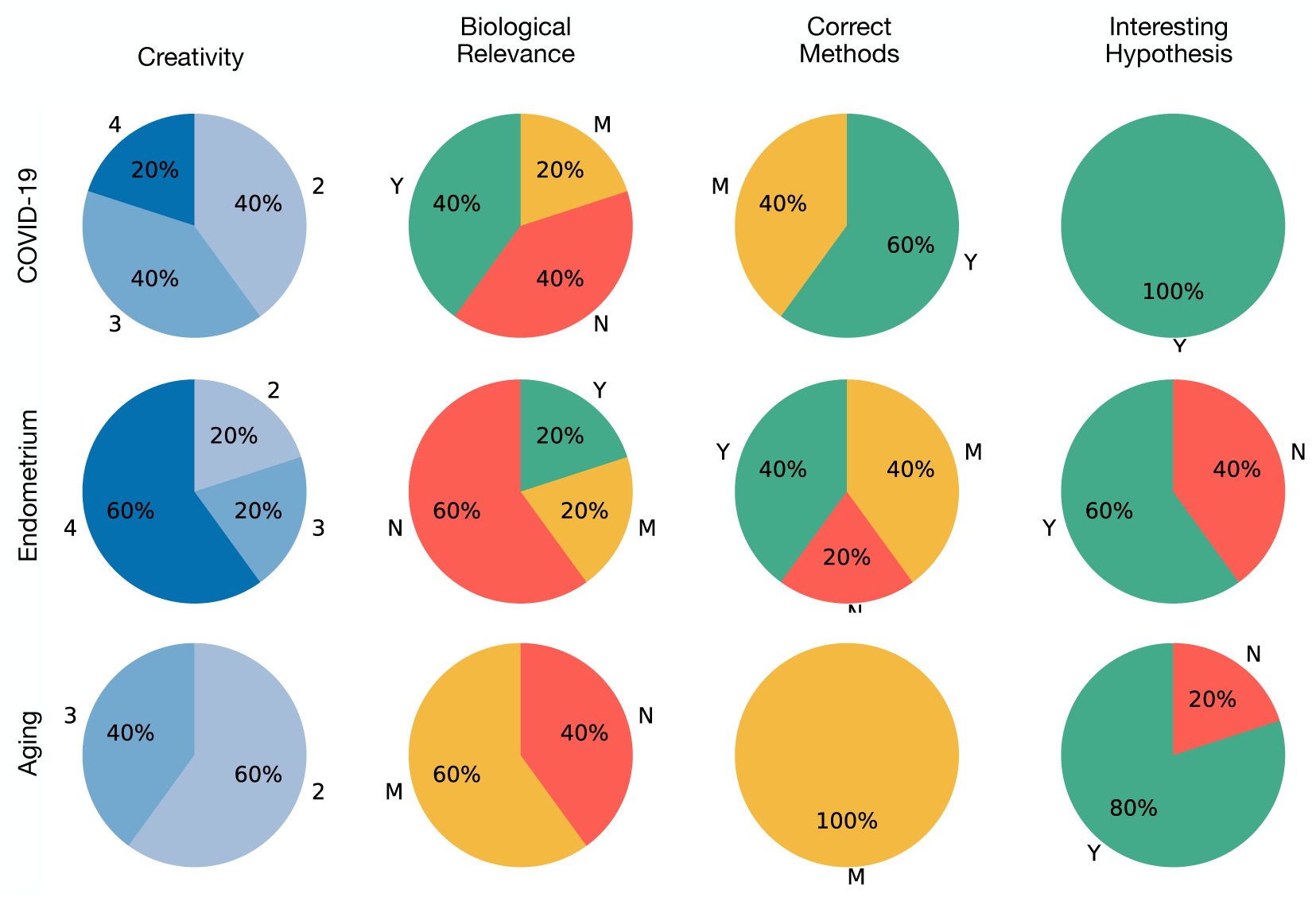
Human expert evaluations. We evaluated CellVoyager on three real-world case studies based on published papers. The authors of these papers evaluated CellVoyager’s feedback on four criteria following a rubric (See **Appendix** Figure 1). For each paper we ran 5 distinct analysis trajectories. Overall, CellVoyager received generally positive feedback in these case studies. Section labels outside the bar correspond to the score or rating given by the evaluators. Creativity was rated on a scale of 1-4. Y: Yes, M: Mostly, N: No.

As a comparison, we attempted to input the same paper information and data files into Google’s Data Science Agent in Colab^1^ after converting them to the required formats. However, the Colab agent failed to produce meaningful results. As a general-purpose data analysis tool operating within Google Colab, it struggled significantly with single-cell data–both in ideation and execution. The analyses it proposed were superficial, and it had difficulty employing domain-specific packages to manipulate the data effectively, often becoming trapped in repetitive error resolution loops to no avail. This observation underscores a broader limitation: generalist data science agents may lack the necessary expertise to handle complex, high-dimensional datasets such as those typical in single-cell analysis. Additionally, this highlights the value of a domain-specific single-cell agent, equipped with deeper biological context, more effective idea generation, and more robust code-writing capabilities.

### 5.1 COVID-19 PBMCs

The first case study is based on a published single-cell transcriptomic atlas of peripheral blood mononuclear cells (PBMCs), which contains samples from seven patients with severe COVID-19 and six healthy controls [49]. The original study found depletion of multiple immune cell subsets (e.g. NK cells and pDCs) and the expansion of plasmablasts, particularly in patients with acute respiratory distress syndrome (ARDS). Furthermore, widespread HLA-class II downregulation was observed in monocytes and B cells, while interferon-stimulated gene signatures were seen to vary by patient age and time since the onset of the fever. Given this rich dataset and the extensive literature on COVID-19 PBMCs [38, 41], there is substantial potential to explore the dataset in new ways.

CellVoyager generated five distinct analyses, each encapsulated in a standalone Jupyter notebook. These analyses were reviewed by one of the original paper’s authors using a standardized rubric (**Appendix Figure 1**). The agent achieved a mean creativity score of 2.8/4 and three of the five analyses were judged to yield biologically meaningful insights. Notably, all five analyses proposed hypotheses that the author thought warranted deeper investigation (**Figure 3**).

Among the most novel analyses was one focused on pyroptosis, a highly inflammatory form of cell death, which received a perfect creativity score (4/4) and was judged to raise biologically meaningful insights (**Figure 4**). Although the inputted paper did not look at pyroptosis, several subsequent publications hypothesized that SARS-CoV-2-induced pyroptosis of monocytes and that macrophages may be an important inflammatory trigger in SARS-CoV-2 infection that may drive the severity of COVID-19 disease [14, 21, 40]. Higher expression of genes involved in pyroptosis may indicate that a cell is more poised to undergo pyroptosis with the appropriate stimulation. CellVoyager began by computing apoptosis and pyroptosis scores based on the expression of pathway-associated genes, comparing these between COVID-19 and healthy samples (**Figure 4A**). Specifically, apoptosis scores were based on the expression of *CASP3*, *CASP8*, *BCL2*, *FADD*, and *TRADD* while pyroptosis scores were based on the expression of *GSDMD*, *CASP1*, *NLRP3*, and *IL18*. CellVoyager first assessed correlations between these cell-level scores and two cell-level gene module scores computed in the original publication: a module of type I IFN-stimulated genes (IFN1) and a module of HLA class II encoding genes (HLA1), both of which correlated with disease severity. This analysis did not reveal a strong correlation between apoptosis or pyroptosis gene scores and IFN1 or HLA1 (**Figure 4B**). The agent next featurized CD14^+^ monocytes using their pyroptosis and HLA1 module scores. When visualizing the resulting groups, it revealed subpopulations enriched for cells from patients with COVID-19, suggesting that these two scores can help isolate COVID-19 CD14^+^ monocytes (**Figure 4C**). Finally, the agent isolated all high pyroptosis CD14^+^ monocytes, split them according to whether they were from COVID-19 or healthy patients, and performed differential gene expression analysis between these two groups (**Figure 4D**). The author of the original paper noted the results in **Figure 4A** could be more statistically rigorous by first pseudobulking each cell type by donor (**Appendix Figure 2**). We provided this feedback into CellVoyager, which successfully completed the analysis and extended it to other celltypes, outputting the visualization seen in **Figure 4E** (some cell types were omitted for readability).

**Figure 4:**
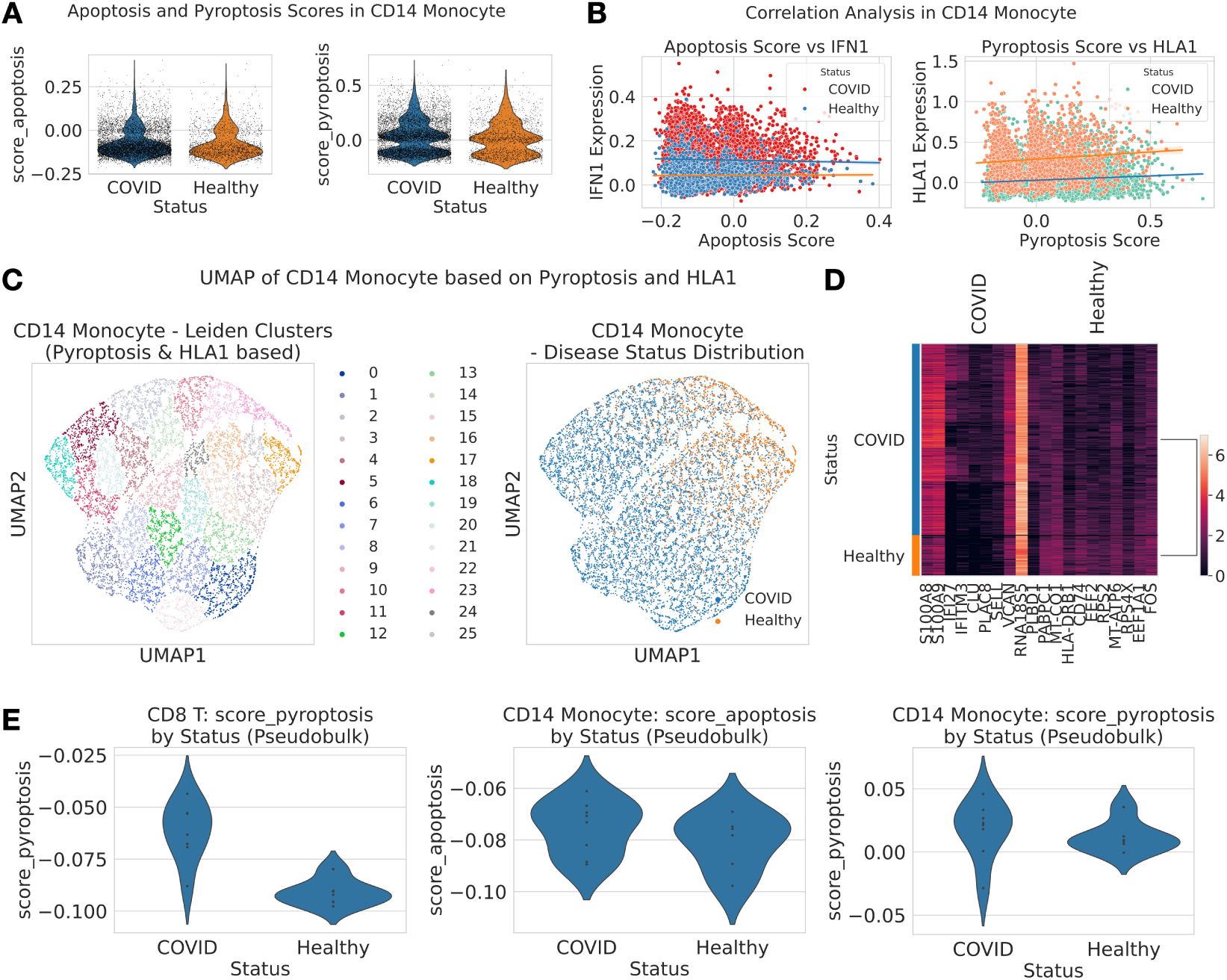
Agent generated analysis from the COVID-19 PBMCs case study. All of the panels are generated by the CellVoyager Agent with some of the figures’ text being altered for readability. **A. Apoptosis and pyroptosis scores**. Apoptosis scores (left) and pyroptosis scores (right) are shown for CD14^+^ monocytes in cells from patients with COVID-19 vs healthy controls. **B. Apoptosis and pyroptosis scores versus module scores.** Apoptosis vs IFN1 scores (left) and pyroptosis vs HLA1 scores (right) are shown for CD14^+^ monocytes. Points are colored by the donor of origin, which is either a patient with COVID-19 or a healthy control. **C. UMAP projection** of CD14^+^ monocytes using pyroptosis and HLA1 scores. Points are colored by their Leiden clusters (left) and their status (right). **D. Heatmap showing differentially expressed genes** between high-pyroptosis scoring monocytes in COVID-19 vs healthy patients **E. Pseudobulked comparison of pyroptosis and apoptosis scores** between COVID-19 samples and healthy samples for CD14^+^ monocytes (middle, right) as well as CD8^+^ T cells (left).

While there were trends towards higher pyroptosis scores in CD14^+^ monocytes from COVID-19 patients, the agent described an overt increase in pyroptosis gene scores in CD8^+^ T cells from COVID-19 patients (p = 0.001). This finding has not been prominently discussed in prior literature, and suggest that CD8 T cells in COVID-19 infection are might be more poised to undergo pyroptosis.

### 5.2 Endometrium Atlas

For our second case study, we applied CellVoyager to a single-cell transcriptomics study of the human endometrium across the menstrual cycle [47]. This study used Fluidigm C1 and 10xGenomics Chromium to profile 73,180 cells from 29 healthy ovum donors sampled across the 28-day menstrual cycle. From this dataset, the original paper identified seven endometrial cell types, including a previously uncharacterized ciliated epithelium, across four major phases of endometrial transformation. Using the same procedure and evaluation protocol as in the previous case study, we sent five CellVoyager-generated analyses to one of the paper’s authors for expert review. Across the five analyses, CellVoyager received a mean creativity score of 3.4/4 and two of the five analyses were judged to yield biologically meaningful insights. Three of the five analyses presented hypotheses that the author found compelling and wanted to investigate further.

**Figure 5** shows an example of an interesting new analysis conducted by CellVoyager, which was not found in the original paper. The agent begins by hypothesizing that “paracrine signaling between stromal fibroblasts and endothelial cells … has distinct correlation patterns in early versus late phases” of the menstrual cycle. Before directly testing this hypothesis, the agent first performs several exploratory analyses. This includes visualizing the distribution of several relevant cell types across different menstrual cycle days (**Figure 5A**) and visualizing stromal fibroblasts using a UMAP, with cells colored by cluster and menstrual cycle day (**Figure 5B**).

**Figure 5:**
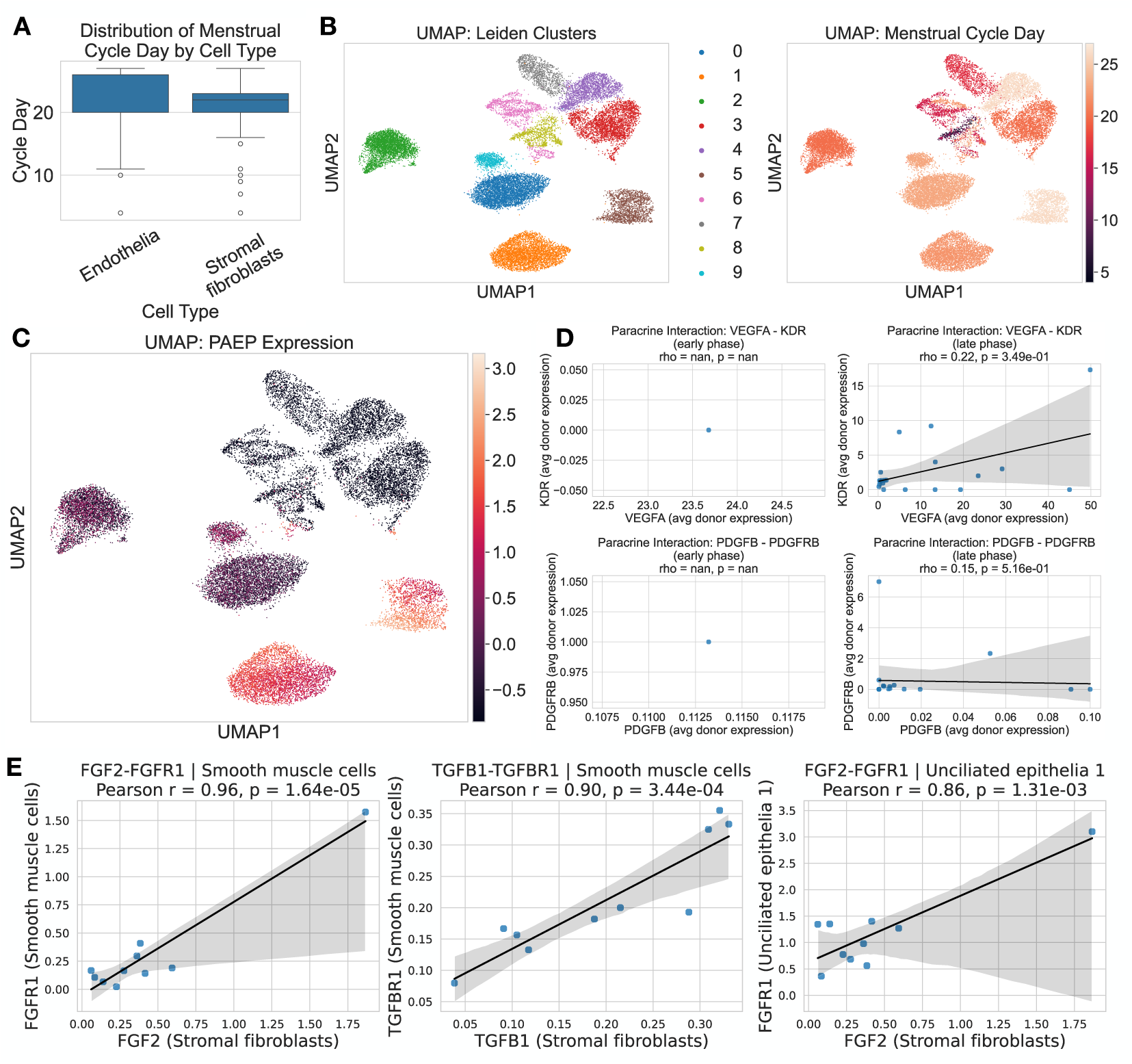
Agent generated analysis for the human endrometrium case study. All the panels are generated by the CellVoyager Agent with some of the figures’ text being modified for readability. **A. Number of cells across cell cycle days** for endothelia and stromal fibroblasts **B. UMAP projection** of the dataset colored by leiden cluster (left) and menstrual cycle day (right) **C. UMAP plot colored by *PAEP*** expression **D. Receptor ligand correlations** between endothelial cells (receptor) and stromal fibroblasts (ligand). On the first row, *VEGFA* vs *KDR* correlations are shown between early phase cells (left) and late phase cells (right). On the second row we have *PDGFB* versus *PDGFBR* in early phase cells (left) and late phase cells (right) **E. Expanded receptor ligand correlations.** These are the most promising receptor-ligand correlations selected by the agent from a set of 40 receptor-ligand pairs the agent examined; the ones selected include *FGF2-FGFR1* and *TGF*β*1-TGF*β*R1* with the former including two different celltypes for expressing the receptor (smooth muscle cells and uncilitated epithelia). As in the previous figure, the ligand is expressed in stromal fibroblasts.

As an illustrative example, it highlights PAEP — a gene emphasized in the original study — revealing high expression in clusters such as 1 and 7 (**Figure 5C**). While not intended as a novel finding, this step demonstrates the agent’s ability to recognize and contextualize known phase-associated gene expression patterns. Finally, to test its hypothesis around paracrine signaling, the agent examined co-expression patterns between receptors from endothelial cells and ligands from stromal fibroblasts. The agent plotted the co-expression of *VEGFA* versus *KDR* and *PDGFB* versus *PDGFRB*, finding positive but statistically insignificant correlations among late phase cells (**Figure 5D**). Notably, there was not enough data among early phase cells to find early phase correlations.

In their review of this analysis, the paper’s author recommended building upon the results in **Figure 5D** by trying to expand the set of ligand-receptor pairs to investigate cell-cell communication (**Appendix Figure 2**). Given this feedback, CellVoyager tested 40 different ligand-receptor correlations. These consisted of five ligand-receptor pairs where the ligand came from stromal fibroblasts and the receptor was tested across 8 different cell types (**Appendix Figure 18**). To facilitate interpretation, we asked the agent to extract and plot the most meaningful correlations, which are shown in **Figure 5E**. TGF-β signaling between fibroblasts and other cell types across the menstrual cycle was among the top pathways identified by the agent followed by the FGF2-FGFR1 signalling across the same cell types. These pathways are recognized as key regulators of cell proliferation and tissue remodeling across multiple systems [33, 53], though the role of these receptor-ligand interactions between stromal fibroblasts and smooth muscle cells, as revealed by the agent, is not fully understood. Their menstrual cycle-dependent activity is consistent with the known roles and may reflect their function in coordinating stromal-epithelial interactions during endometrial regeneration.

### 5.3 Aging in the Brain

Our last case study builds off a paper where single-cell RNA-seq data was generated from the subventricular zone of adult mouse brain collected across multiple ages [7]. The study identified 11 cell types, including cell types of the neural stem cell lineage, and built single-cell aging clocks, machine learning models trained to predict age, for six of these cell types. The original dataset was used to train these aging clocks, which were then deployed to profile the effect of various rejuvenation interventions such as exercise and heterochronic parabiosis on the aging of different cell types. The original dataset was also leveraged to understand age-related gene expression trajectories and gene pathways that contributed to age prediction in the aging clocks. Given the targeted focus of the original study on building and applying aging clocks, there is substantial opportunity for new insights in re-analysis of this resource.

We again input the paper text as well as the processed single-cell dataset into CellVoyager and sent agent-generated analyses to an author on the paper. Across the five analyses, the agent received a mean creativity score of 2.4 with three of the analyses yielding some biological insight and four of the analyses raising an interesting hypothesis that the author wanted to investigate further (Table 1). An example among these was the hypothesis that “transcriptional noise in Astrocyte_qNSC cells increases with age, reflecting a loss of regulatory precision associated with cellular aging in the brain’s neurogenic niche” (**Figure 6**). Transcriptional noise was not analyzed in the original study, but increasing transcriptional noise has been linked to aging through several recent studies [4, 36, 44]. Understanding whether such transcriptional noise also increases in astrocytes and quiescent neural stem cells (qNSCs) during aging is of interest in the adult mouse subventricular zone given the role that these cells play in neurogenesis, which declines with age [43].

**Figure 6:**
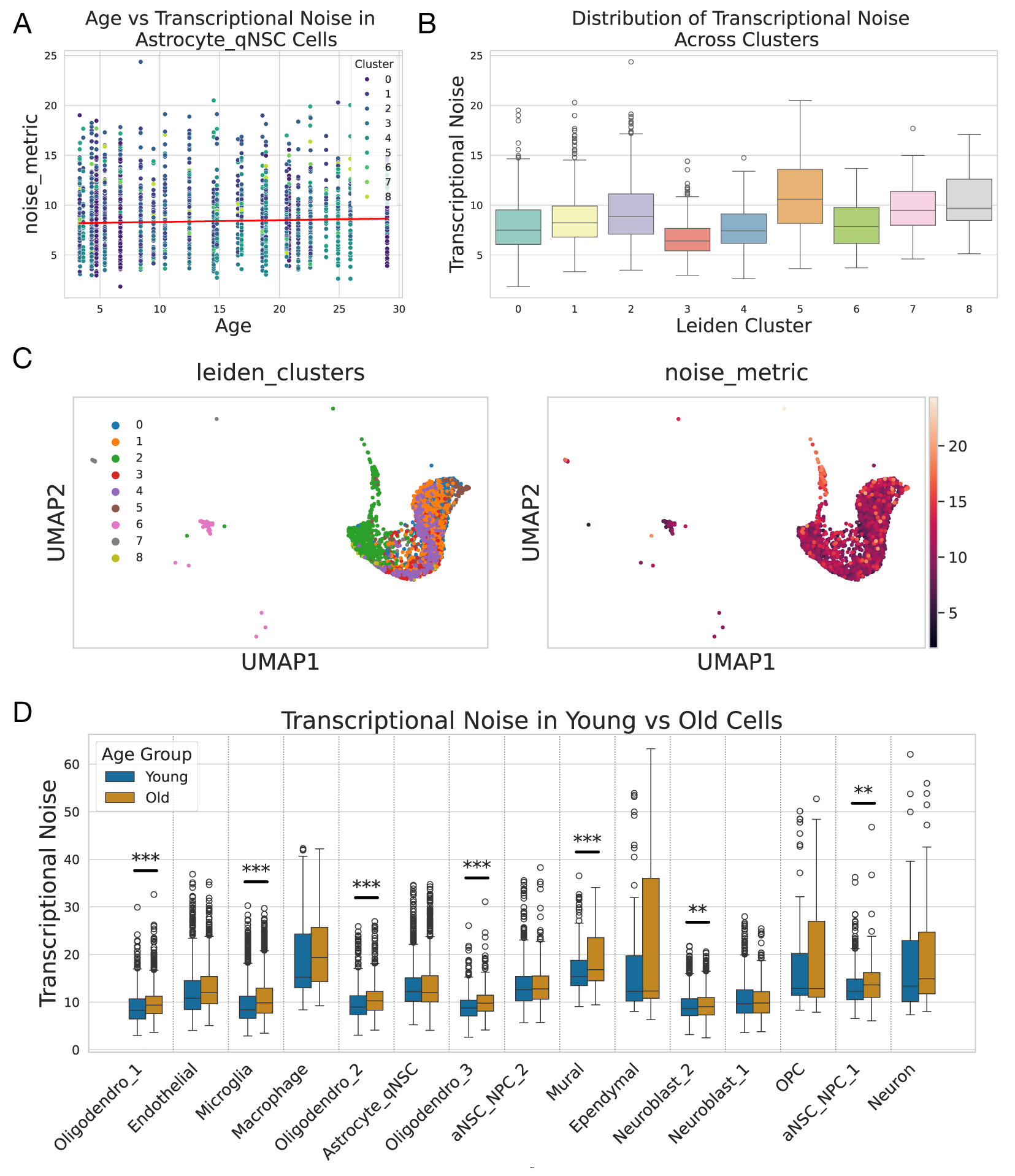
Agent generated analysis for the aging in the brain paper. All the panels are generated by the CellVoyager Agent with some of the figures’ text being modified for readability. **A. Transcriptional noise versus age** in the astrocyte-qNSC cluster. Points are colored by their Leiden cluster. **B. Distribution of transcriptional noise** per Leiden cluster in astrocyte-qNSC cells. **C. UMAP projection of astrocyte-qNSC cells** colored by their respective Leiden clusters (left) and their transcriptional noise (right). **D. Transcriptional noise in young versus old cells** for all of the celltypes in the dataset. For statistical significance, ^∗^: *p* < 0.05,^∗∗^: *p* < 0.01,^∗∗∗^: *p* < 0.001

The analysis performed by the agent begins with standard preprocessing of single-cell RNA-seq count data, including normalization and log-transform with a pseudocount. The agent then subsets the astrocyte-qNSC cluster containing the cell type of interest and subclusters these cells further using Leiden clustering, before measuring transcriptional noise using Euclidean distance to the subcluster centroid using the top ten principal components. The approach used by the agent for measuring transcriptional noise is similar to that from previous single-cell aging studies [1, 13, 23], with a notable difference being an additional subclustering step before computing distances. The agent reported a marginal positive correlation between the transcriptional noise and age across all subclusters (r = 0.047, *p* = 0.0135), suggesting that there is relatively little change in transcriptional noise of Astrocytes-qNSCs in the subventricular zone during aging (**Figure 6A**). The observation is consistent with findings with meta-studies indicating that transcriptional noise may not be a universal hallmark of aging [19]. Interestingly, the agent also discovered that the transcriptional noise varied between different subclusters of Astrocytes-qNSCs, suggesting that some subpopulations may be more susceptible to these changes during aging (**Figure 6B**).

After reviewing the initial results, the original study author asked CellVoyager to visualize the Leiden clusters referenced in **Figure 6B** and to color them by their transcriptional noise (**Appendix Figure 2**). CellVoyager successfully executed this request, producing the plots shown in **Figure 6C**. Seeing little signal for Astrocytes-qNSCs, the author then asked CellVoyager to extend its analysis across all celltypes, prompting the agent to look at young versus old cells (split by the median age) rather than examining correlations (**Appendix Figure 2**). The agent’s results in **Figure 6D** demonstrate a general increase in transcriptional noise for older cells relative to younger cells across all cell types. In particular, oligodendrocytes (across all 3 subclusters), microglia, and mural cells had significantly higher transcriptional noise in older cells versus younger cells (*p* < 0.001). Although transcriptional noise has been reported to increase with age for these cell types in multiple brain regions [54], the agent’s analysis provides the first indication that these age-related changes are also conserved in the subventricular zone neurogenic niche.

## 6 Discussion

CellVoyager is an LLM-based agent capable of autonomously exploring scRNA-seq data and complementing prior work done by a researcher. To test the latter capability in a challenging setting, we focus on how well CellVoyager can expand upon the results presented in a published scRNA-seq research paper, where extensive effort has been put into fully exploring the dataset. Using CellBench, we evaluated CellVoyager’s ability to propose analysis ideas that are consistent with the kinds of analyses typically undertaken in single-cell studies. Our results suggest that CellVoyager can generate analysis ideas that are appropriate given a research direction, outperforming the state of the art LLMs in this task.

To assess CellVoyager’s performance in a research setting, we conducted three case studies using published manuscripts on single-cell analysis and their associated datasets. In each case, we evaluated whether the agent could propose and execute creative, biologically meaningful analyses distinct from those included in the original papers. Authors of the original studies, all of whom hold PhDs relevant to this domain, rated each analysis using a standardized rubric. Despite the difficulty in expanding upon a research paper, CellVoyager was able to propose creative analyses distinct from those tried in the paper (average score of 2.8/4) with 12 of the 15 total proposed hypotheses being judged to be interesting enough to warrant further investigation. CellVoyager was also able to find several new biological insights, such as higher pyroptosis scores in CD8+ T cells in COVID-19 patients relative to healthy patients. Nevertheless, the authors noted that several of the analyses executed by CellVoyager could be improved with a few additional steps and/or corrections. Therefore, we integrated human-in-the-loop capabilities for CellVoyager where the authors’ comments can be given back to the agent. We demonstrated that the agent was able to meaningfully improve its analyses using just one to two pieces of feedback from the author (See **Section 5**). This suggests that although CellVoyager does well at finding creative and insightful analysis trajectories, it still benefits from user feedback to help guide it towards more biologically meaningful conclusions. This capability further enhances CellVoyager’s ability to act as a collaborative tool for researchers, who can combine their expertise with CellVoyager’s exploratory capabilities to find more biological insights in their data.

While CellVoyager is capable of proposing and executing meaningful analysis on single-cell data, our work has several limitations. In the current version, CellVoyager runs analyses sequentially, taking around 30 minutes to run each analysis. One possible addition to speed up run time is to identify independent lines of analysis and performing them in parallel. However, in this work, we focus more on the analytical capabilities of the agent instead of minimizing run time. This work also opens up numerous future additions. Currently, we focus on single-cell transcriptomic data, but it is by no means the only domain in biology with high dimensional data that require domain-specific tools and knowledge to analyze. The design of CellVoyager is deliberately modular and flexible to a variety of tasks (see **Figure 1**), though testing on other domains was out of the scope this study. Future work can include generalizing to other domains, such as spatial transcriptomics, proteomics, etc. In our initial experiments, we found that incorporating additional biological contextual information helped the agent propose more relevant analyses. Augmenting CellVoyager with more extensive literature search can potentially boost its performance. Additionally, in our experiments CellVoyager often uses popular Python packages such as Scanpy. Without modification, the agent may be less able to use less popular or custom / in-house tools effectively. While this could be mitigated through finetuning, a potentially efficient method could be to provide tool metadata to the agent to inform its tool use [31]. Lastly, while we designed CellVoyager to be training-free and use LLMs out of the box, future work can investigate how to finetune modules within CellVoyager.

While CellVoyager can help users to analyze new data, our study also demonstrates that many new insights could be gained from reanalysis of existing public data. With over 100,000 published papers in single-cell sequencing alone, thorough reanalysis by human researchers is prohibitive, but agents like CellVoyager could potentially unlock a treasure trove of new insights by tirelessly analyzing these vast public datasets.

## Data Availability

Published papers used in the case studies can be found at https://doi.org/10.1038/s41591-020-0944-y for the COVID-19 study, https://doi.org/10.1038/s43587-022-00335-4 for the aging study, and https://doi.org/10.1038/s41591-020-1040-z for the endometrium study. The associated datasets can be found in the respective publications as well. The CellBench evaluation dataset is available at https://github.com/zou-group/CellVoyager.

## Code Availability

Code for running the agent on case studies and for CellBench evaluation is available in the CellVoyager GitHub repository: https://github.com/zou-group/CellVoyager.

## Acknowledgements

We thank Catherine Blish for helpful feedback during the project. J.Z. is supported by funding from the Chan-Zuckerberg Biohub. This work was supported by the NSF Graduate Research Fellowship under Grant #DGE-2146755 (S.A.).

## A Extended Methods

### A.1 CellVoyager Implementation

Implementation of CellVoyager used Python (v3.9.22) and the following Python packages: reportlab (v4.4.0), nbformat (v5.10.4), and openai (v1.77.0). CellVoyager’s own analyses used the following Python packages: scanpy (v1.10.3), scvi-tools (v1.1.6), celltypist (v1.6.3), anndata (v0.10.8), matplotlib (v3.9.4), numpy (v1.26.4), seaborn (v0.13.2), pandas (v2.2.3), and scipy (v1.13.1).

An overview of the implementation of CellVoyager is highlighted in **Algorithm 1** and **Figure 1**. In the following sections, we go into more detail on each step of CellVoyager’s procedure and showcase the prompts used for those steps.

**Algorithm 1.**
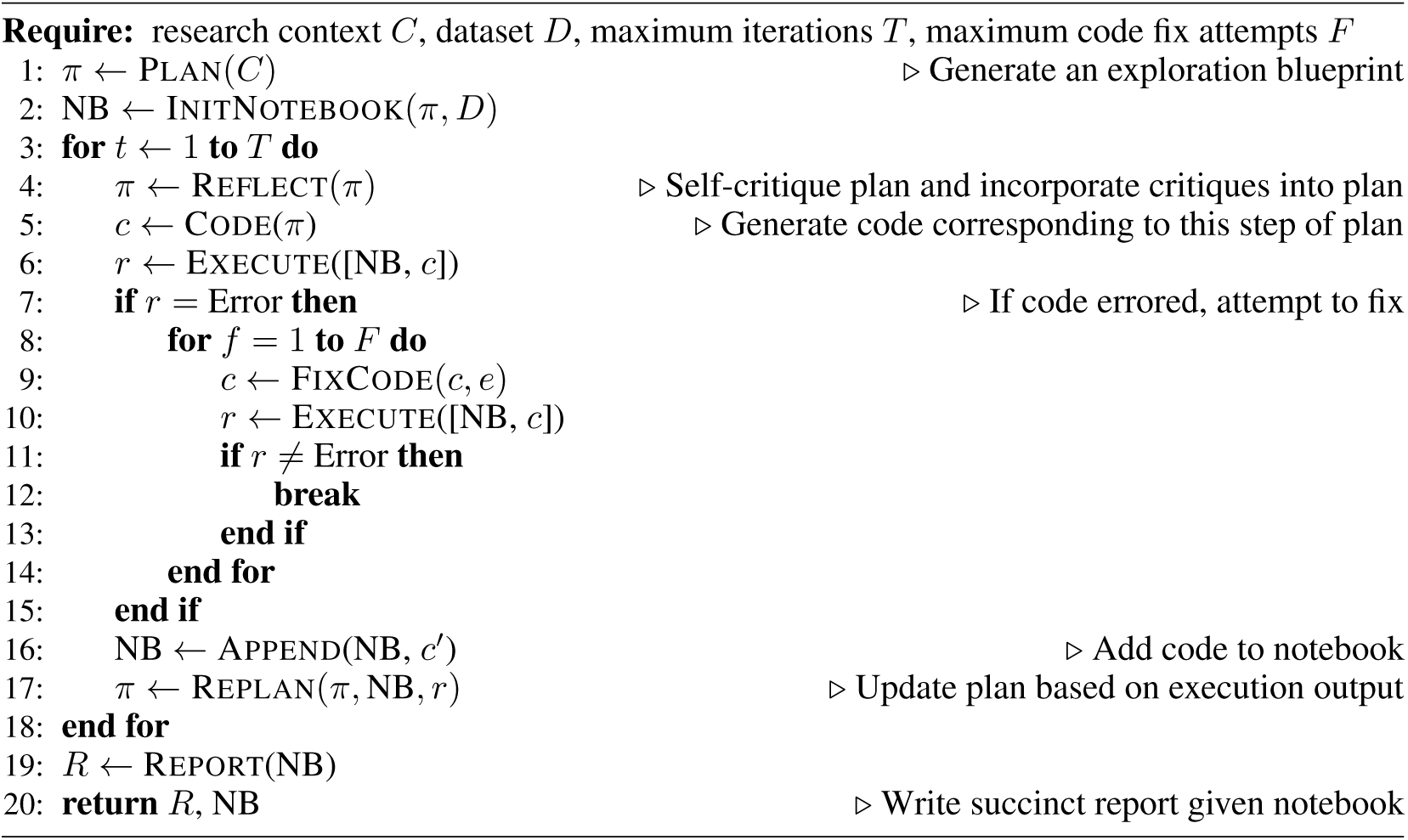
CellVoyager Analysis for a Single Trajectory.

#### A.1.1 Preprocessing

The agent loads in the metadata of the single-cell data object and summarizes it by listing column names and the first 10 unique values from that column (called adata_summary in the prompts). Then, the agent loads in a manuscript or other text files about the biological problem or background and, optionally, a set of analyses already performed on the single-cell dataset. This information is summarized via the prompt shown in **Appendix Figure 3** and is passed into the prompts as paper_txt. The names of Python packages available in a user-configured Python environment are passed into the agent. In this work, we focus on using scanpy, scvi-tools, and anndata since they are popular and compatible with one another. A deep-research style summary of general scRNA-seq analyses compatible with the packages in our environment was inputted into the agent as deepresearch_summary. Finally, the agent initializes a Jupyter notebook by loading in the single-cell dataset and importing all of the necessary packages. Since the Jupyter kernel maintains state, the agent can iteratively run analyses without having to rerun previous analyses at each step.

#### A.1.2 Initial Analysis Plan Generation

CellVoyager is designed to explore datasets as thoroughly as possible, beginning with an empty summary of prior analyses. The first idea for the analysis plan is generated via the prompt shown in **Appendix Figure 4**, which uses the system prompt shown in **Appendix Figure 5**. This process yields a hypothesis, a step-by-step analysis plan, Python code for the first step, a description of that code, and a brief 1–2 sentence summary of the analysis. The agent is also given a set of coding guidelines, which consists of common pitfalls observed when first designing the agent (**Appendix Figure 6**). The agent reflects on its plan by first acting as a critic of the analysis plan (**Appendix Figure 7**) and then incorporating its critiques back into the analysis plan (**Appendix Figure 8**). Although the system is designed to support multiple rounds of reflection, in this work we limit it to a single iteration to reduce runtime, as additional iterations did not yield significant improvements. Finally, the agent adds the initial hypothesis and the full analysis plan to the Jupyter notebook.

#### A.1.3 Executing the Analysis Plan

The agent first adds a markdown cell to the Jupyter notebook that describes the purpose of the code for the first analysis step, followed by a code cell containing the corresponding code. It then attempts to execute the code cell. If execution fails, the agent invokes the prompt in **Appendix Figure 9** to iteratively debug and revise the code, with a maximum of three attempts. Upon successful execution, the textual and visual outputs of the code are passed into a vision language model via the prompt in **Appendix Figure 10**, which produces an interpretation of the results. This interpretation is appended to the notebook as a markdown cell. If the code cannot be successfully fixed after three attempts, the agent instead records a markdown cell noting that the code failed to run and that a different analytical approach should be considered.

#### A.1.4 Re-evaluating the Analysis Plan

Following the execution and interpretation of each step in the analysis plan, the agent updates its strategy based on the observed results. This is achieved using the prompt shown in **Appendix Figure 11** to generate a revised plan starting from the second step of the prior analysis (since the first step has already executed). CellVoyager then proceeds through the same iterative process described above. In this work, we limited the agent to a maximum of eight steps per analysis for efficiency. Upon completion, the agent appends the summary portion of the analysis plan to a list tracking its analysis history (inputted into the prompts as past_analyses), which is used as input when planning the next analyses.

### A.2 CellBench

We searched for published papers that only focus primarily on analyzing a scRNA-seq dataset. For 50 such papers, we used gemini-2.5-pro-preview to extract the biological background and each scRNA-seq computational analysis performed in the paper using the prompt in **Appendix Figure 12**. A subset of the curated examples were reviewed by experts to check that the extracted information accurately reflects the content of the paper.

For inference on CellBench (GPT-4o and o3-mini), we used the OpenAI API with default hyperparameters (e.g., temperature of 1 for GPT-4o, reasoning effort of medium for o3-mini, etc.). See **Appendix Figure 13** for inference prompts used by the base LLMs. To run CellVoyager on CellBench, we adjusted the prompts to match the task of CellBench, i.e., analysis prediction without any coding. In summary, identical to the typical CellVoyager workflow, the agent makes an initial plan (prompt shown in **Appendix Figure 15**), self-reflects to get feedback (**Appendix Figure 16**), and incorporates its feedback (**Appendix Figure 17**) to output its final analysis proposal.

For evaluation, we use LLM-as-a-judge to check if a proposed analysis is represented in the held-out set of original analyses. The LLM judge was verified by human researchers on a subset of the outputs (**Section 4**). The judge prompt is shown in **Appendix Figure 14**.

**Appendix Figure 1:**
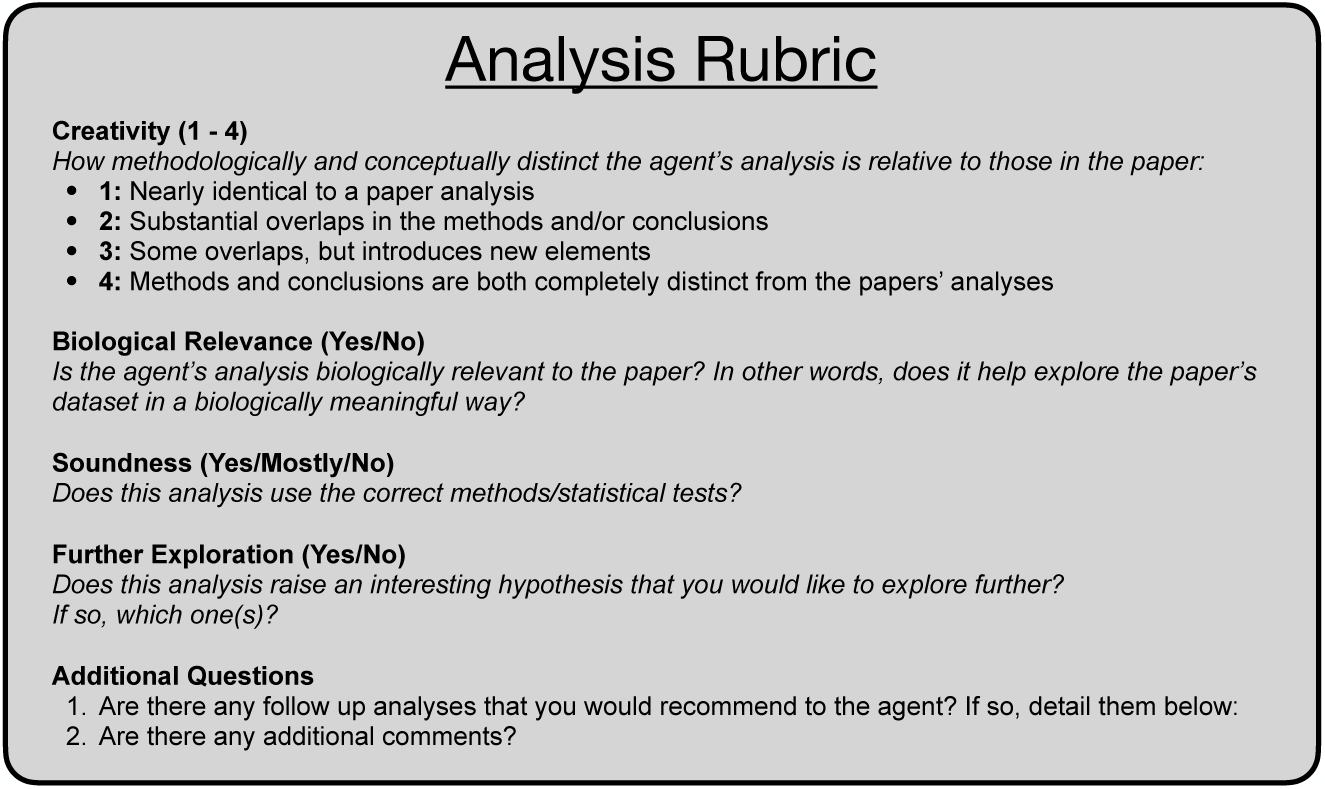
Rubric used for case studies to grade CellVoyager’s analyses.

**Appendix Figure 2:**
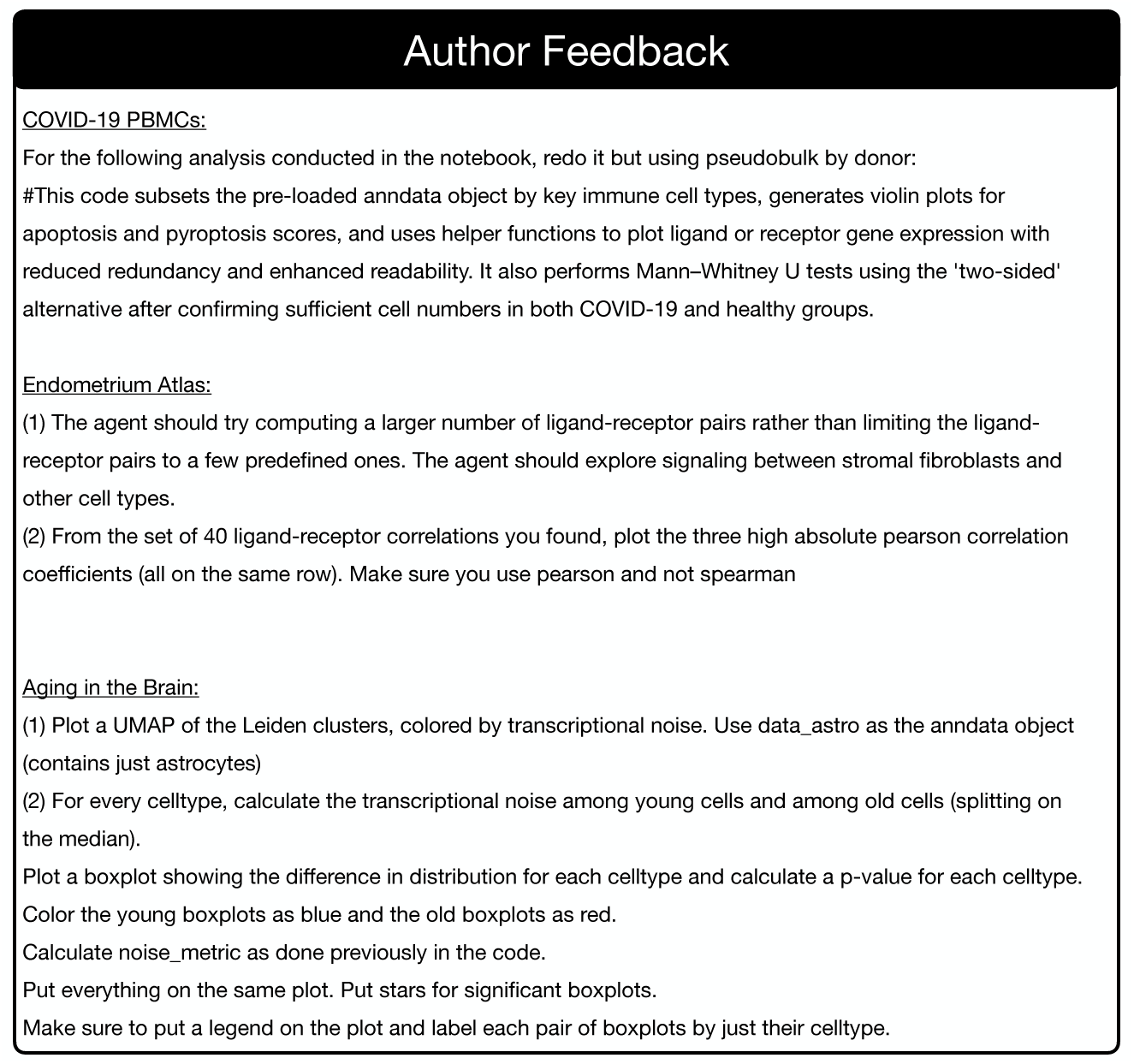
Feedback used from the authors to improve the case study analyses

**Appendix Figure 3:**
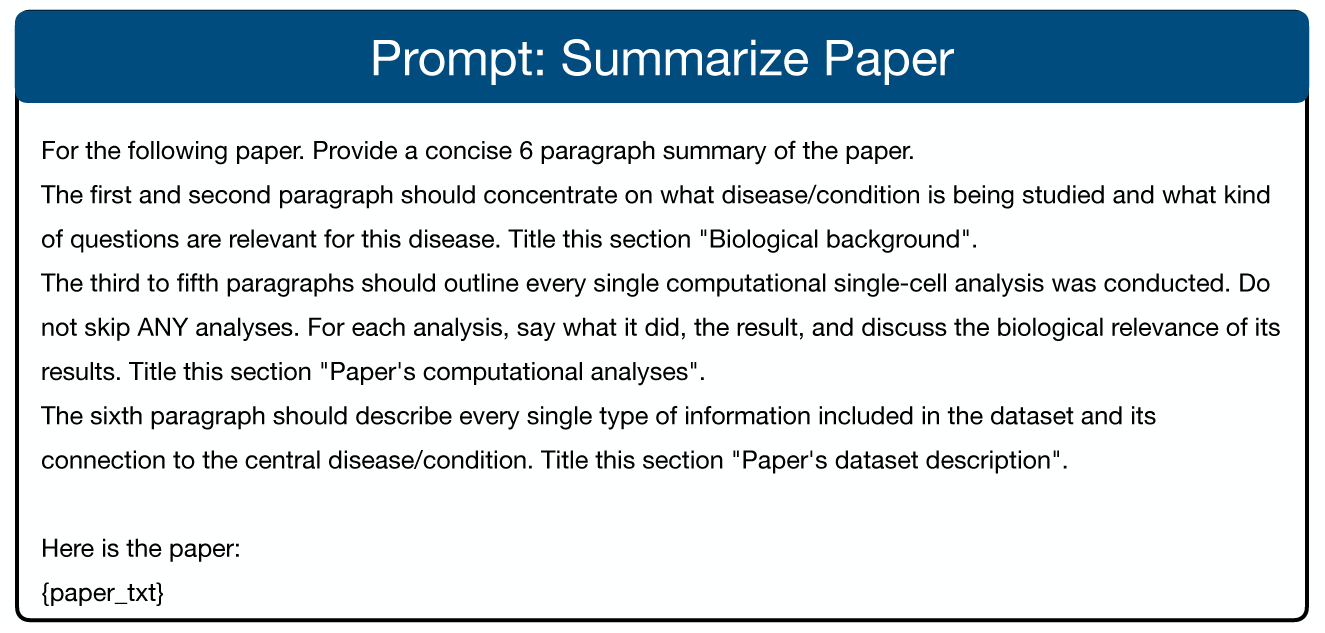
Prompt used for summarizing the inputted manuscript

**Appendix Figure 4:**
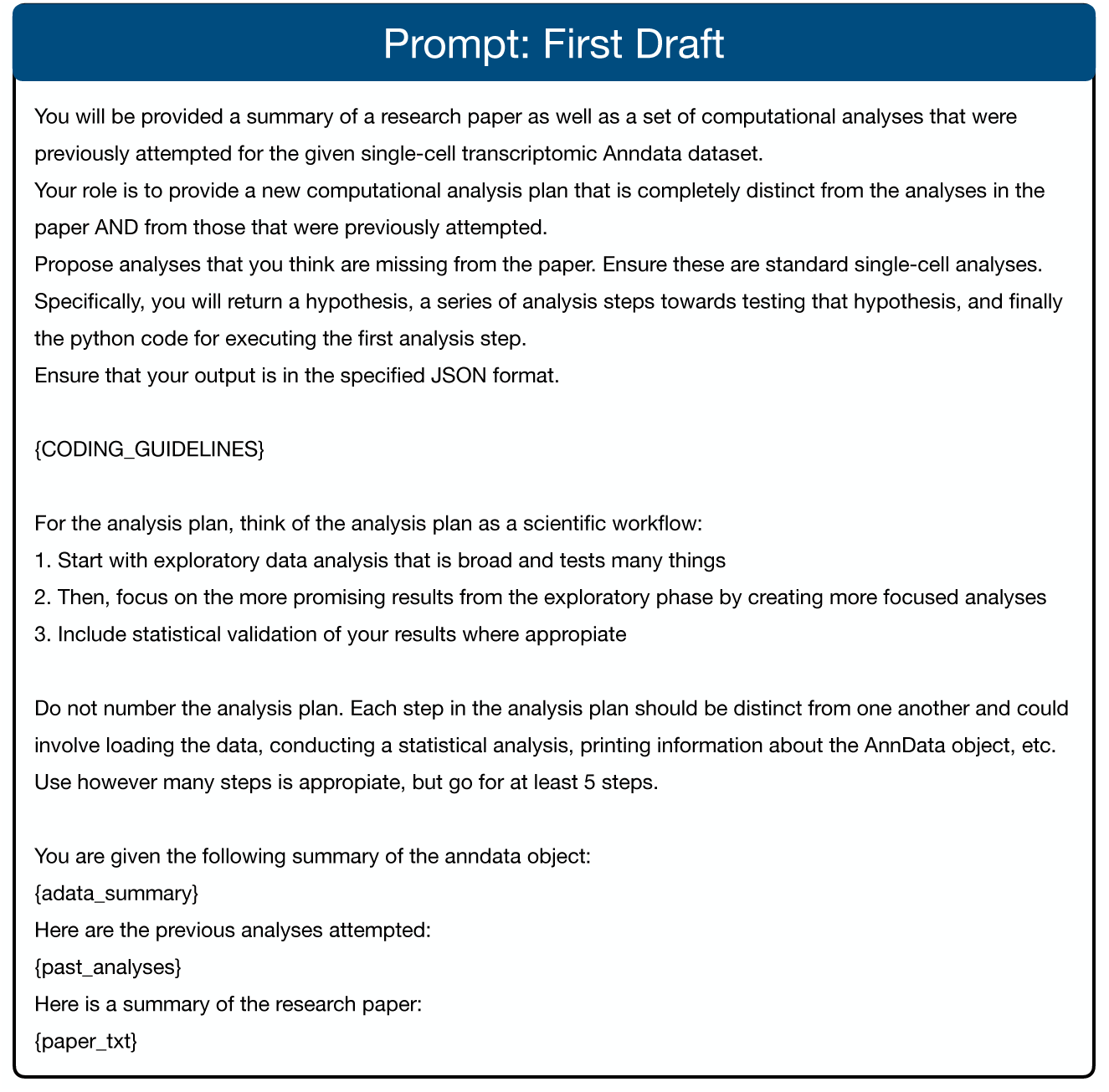
Prompt for generating the first draft of the analysis

**Appendix Figure 5:**
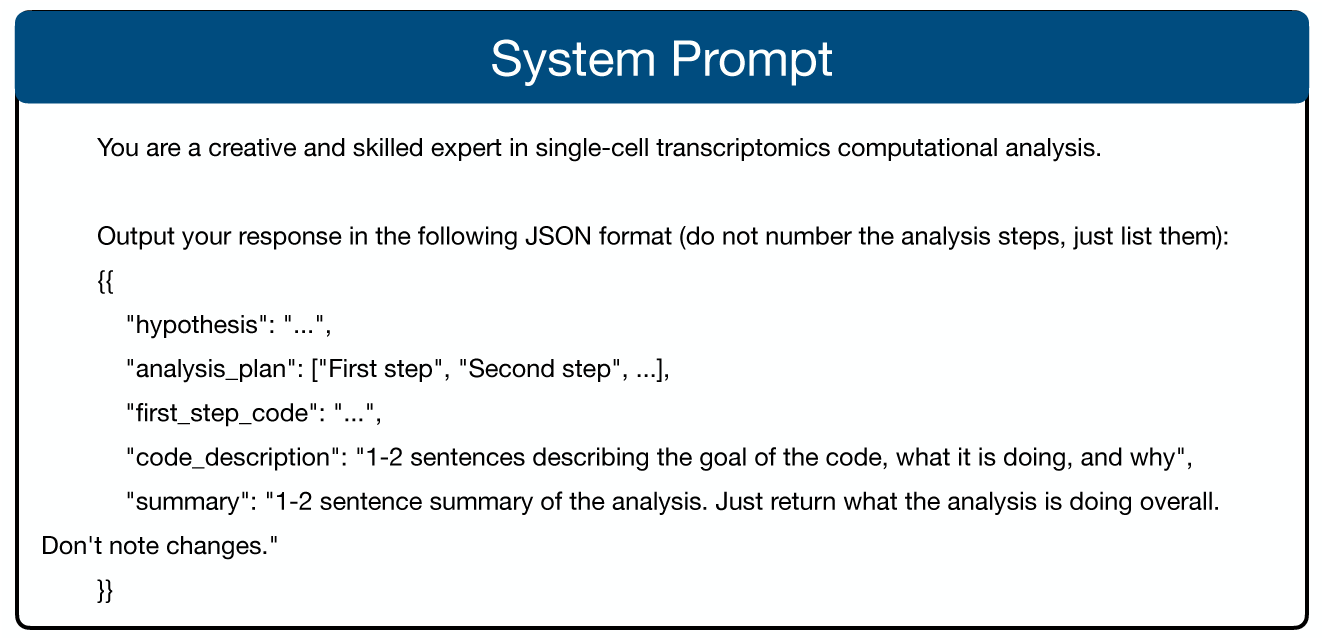
System prompt inputted for all parts of the agent besides fixing the code

**Appendix Figure 6:**
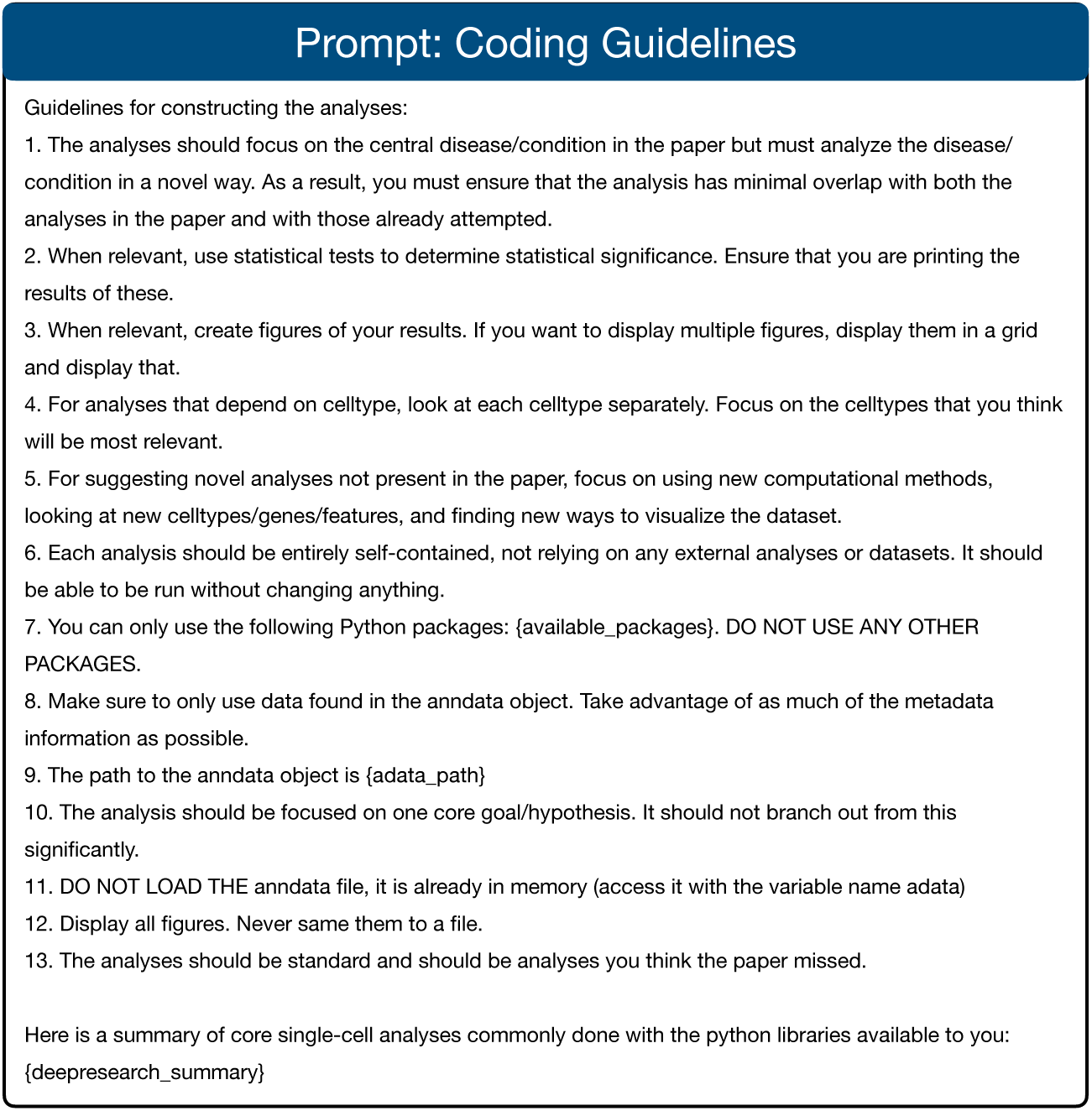
Prompt used for guiding the code generation

**Appendix Figure 7:**
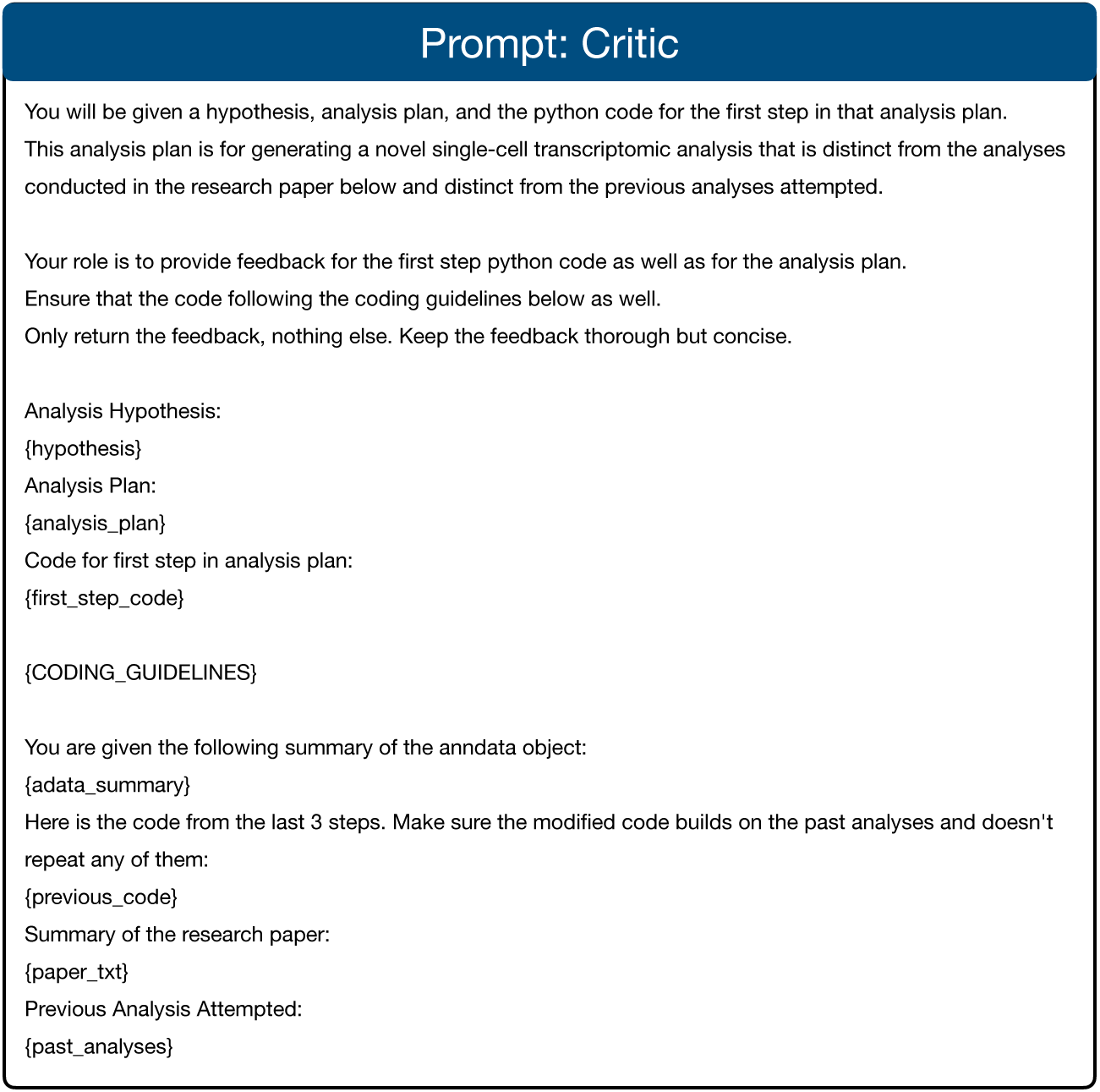
Prompt for the critiquing the analysis plan and code.

**Appendix Figure 8:**
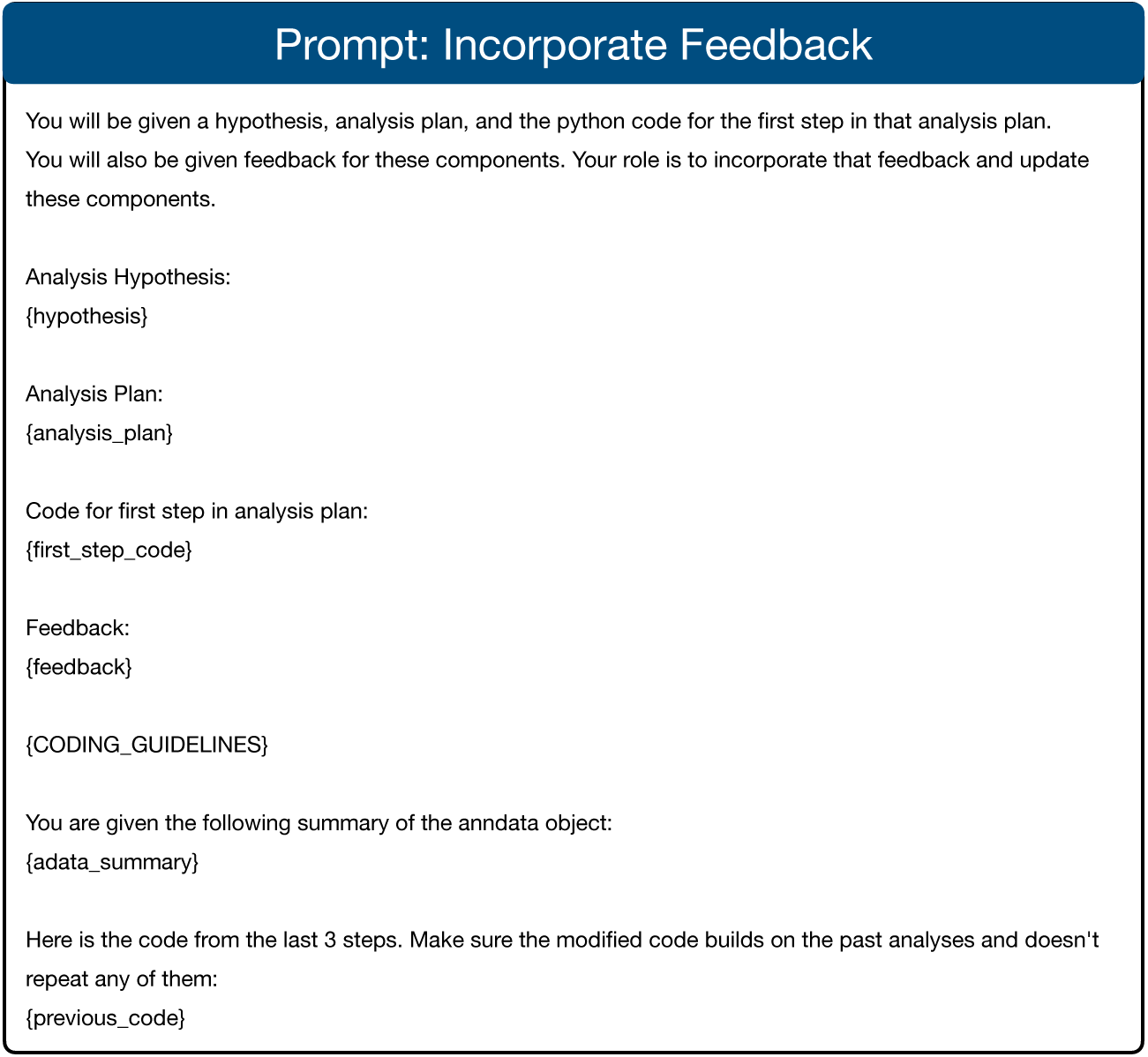
Prompt for incorporating feedback for the analysis plan.

**Appendix Figure 9:**
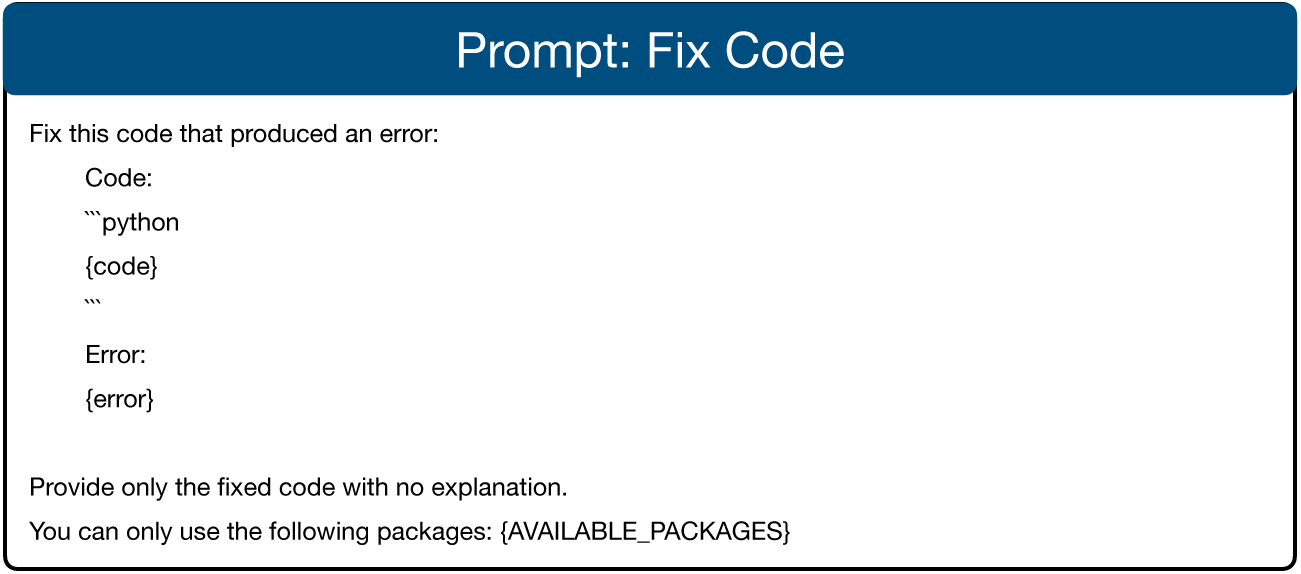
Prompt for fixing an error in the code

**Appendix Figure 10:**
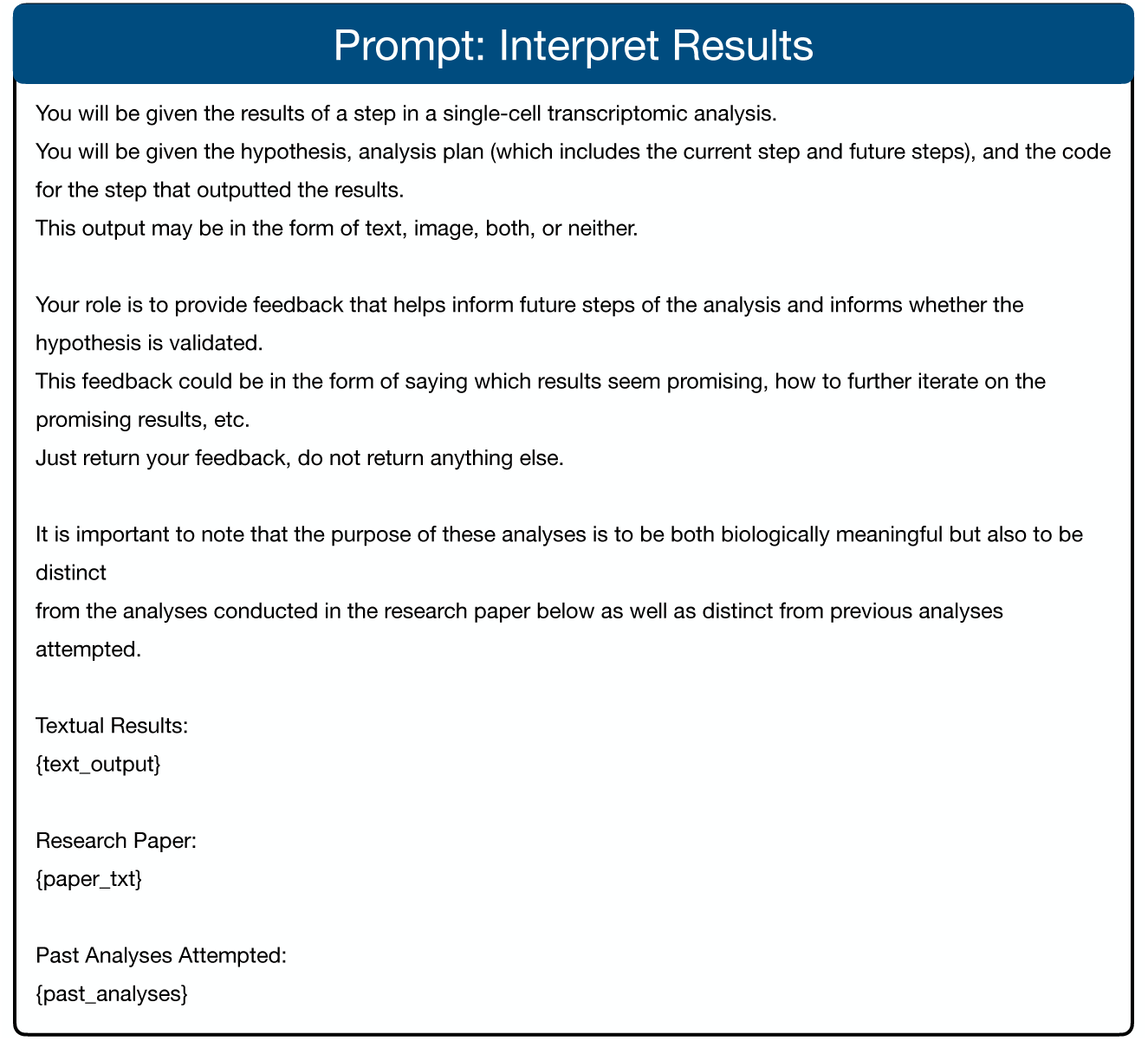
Prompt for interpreting the output(s) from a Jupyter notebook code cell.

**Appendix Figure 11:**
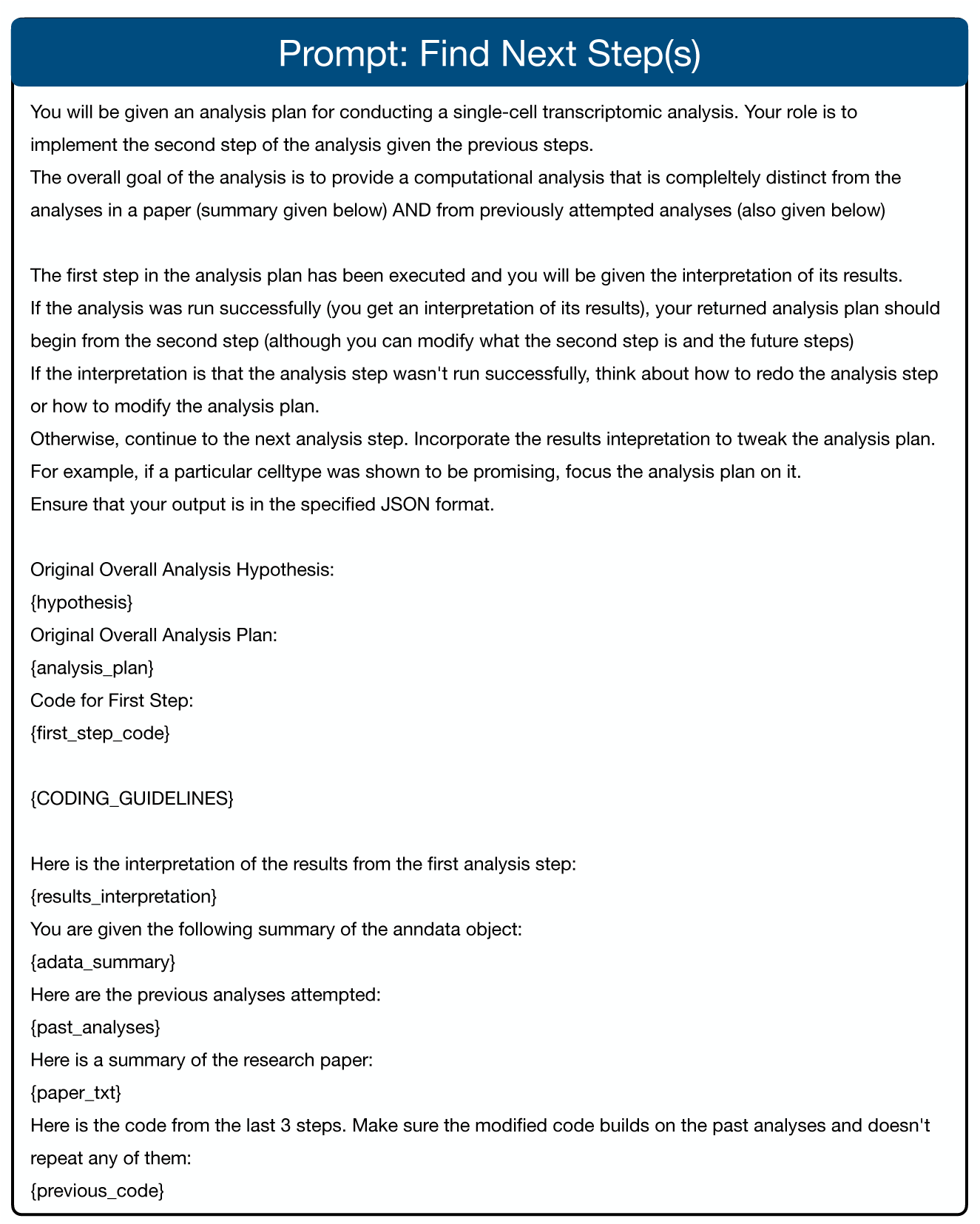
Prompt for generating the next step of the analysis plan, which may involve changing the overall analysis plan depending on the results of previous steps.

**Appendix Figure 12:**
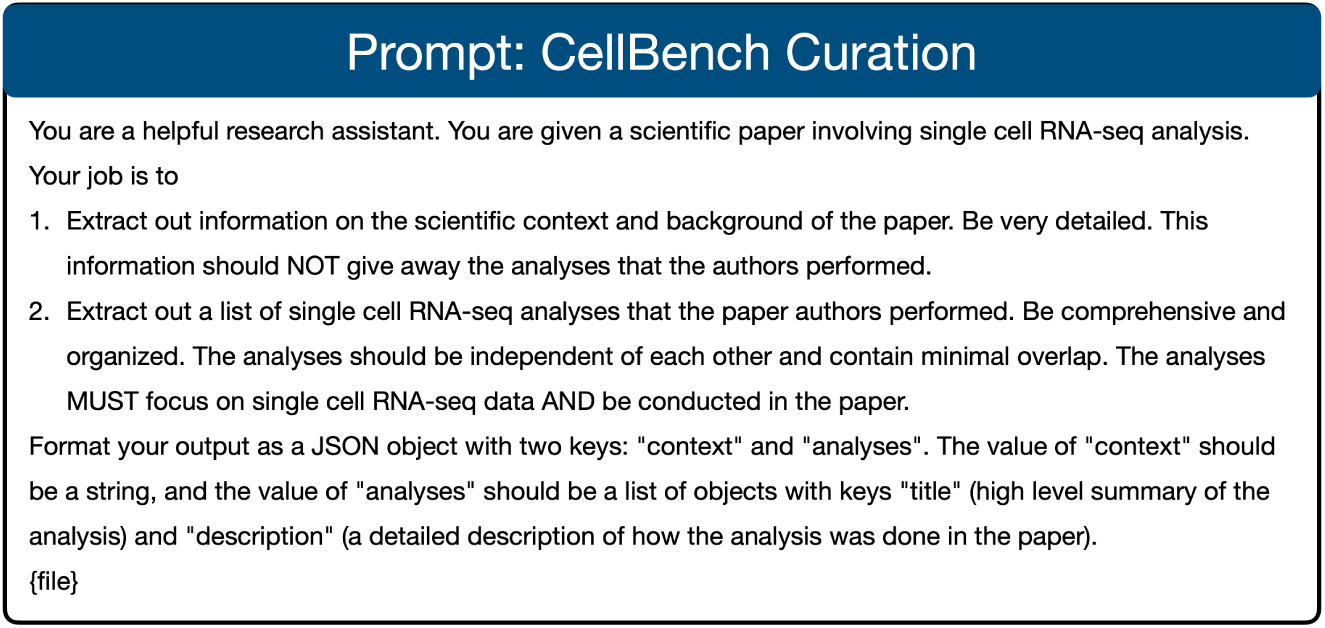
Prompt for extracting background and performed single-cell analyses from papers, used to curate CellBench.

**Appendix Figure 13:**
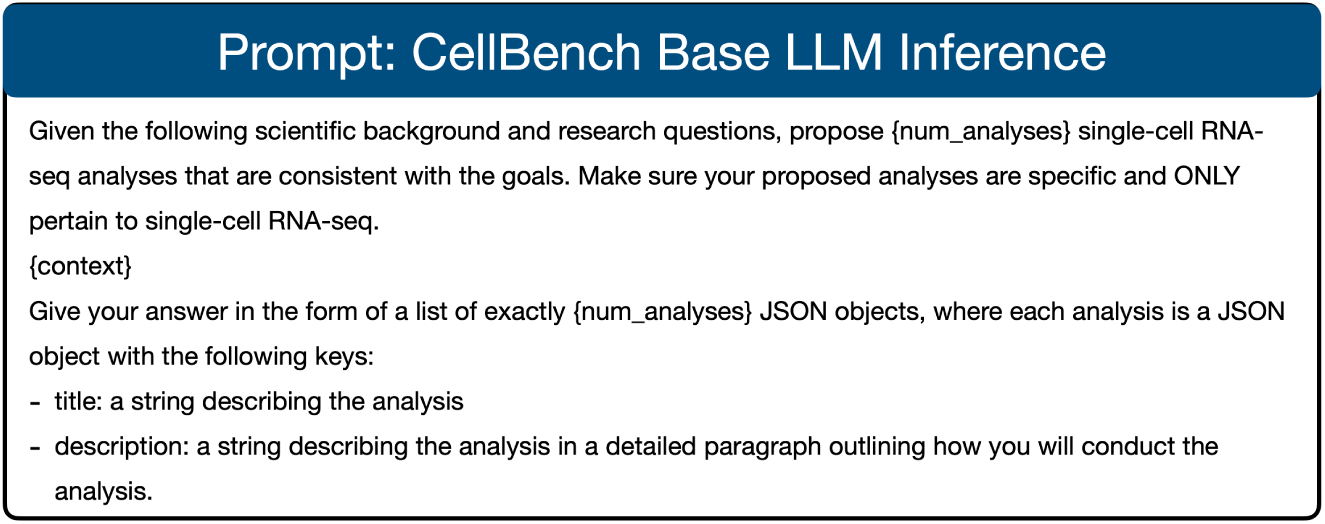
Inference prompt for running base LLMs on CellBench.

**Appendix Figure 14:**
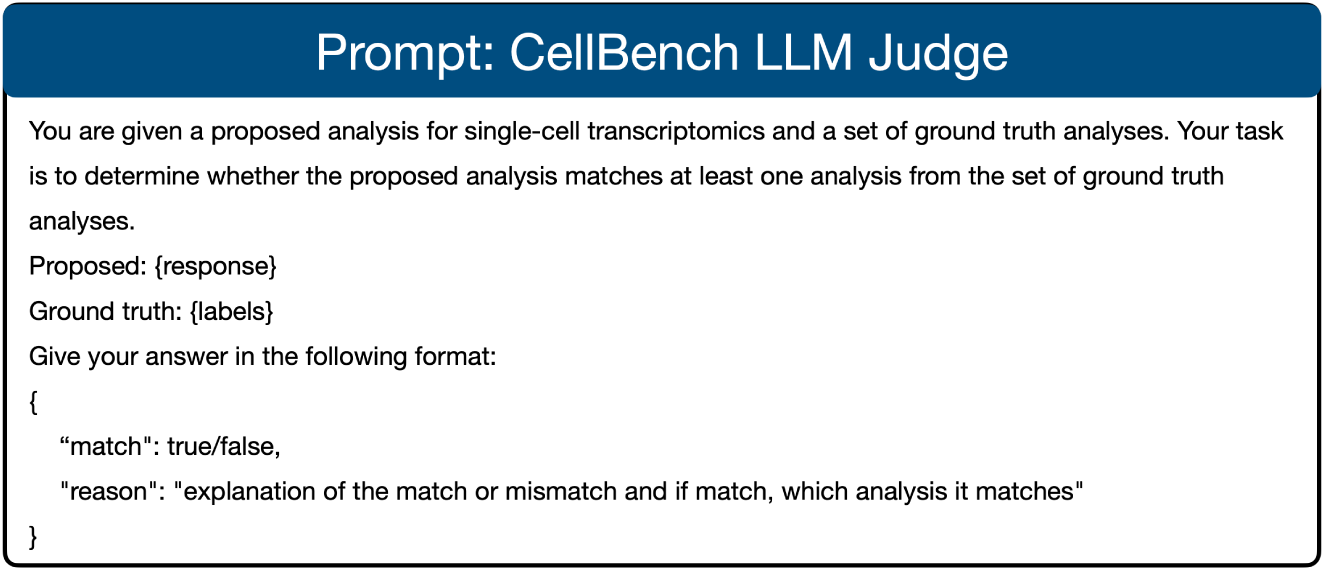
LLM judge prompts used to evaluate CellBench responses.

**Appendix Figure 15:**
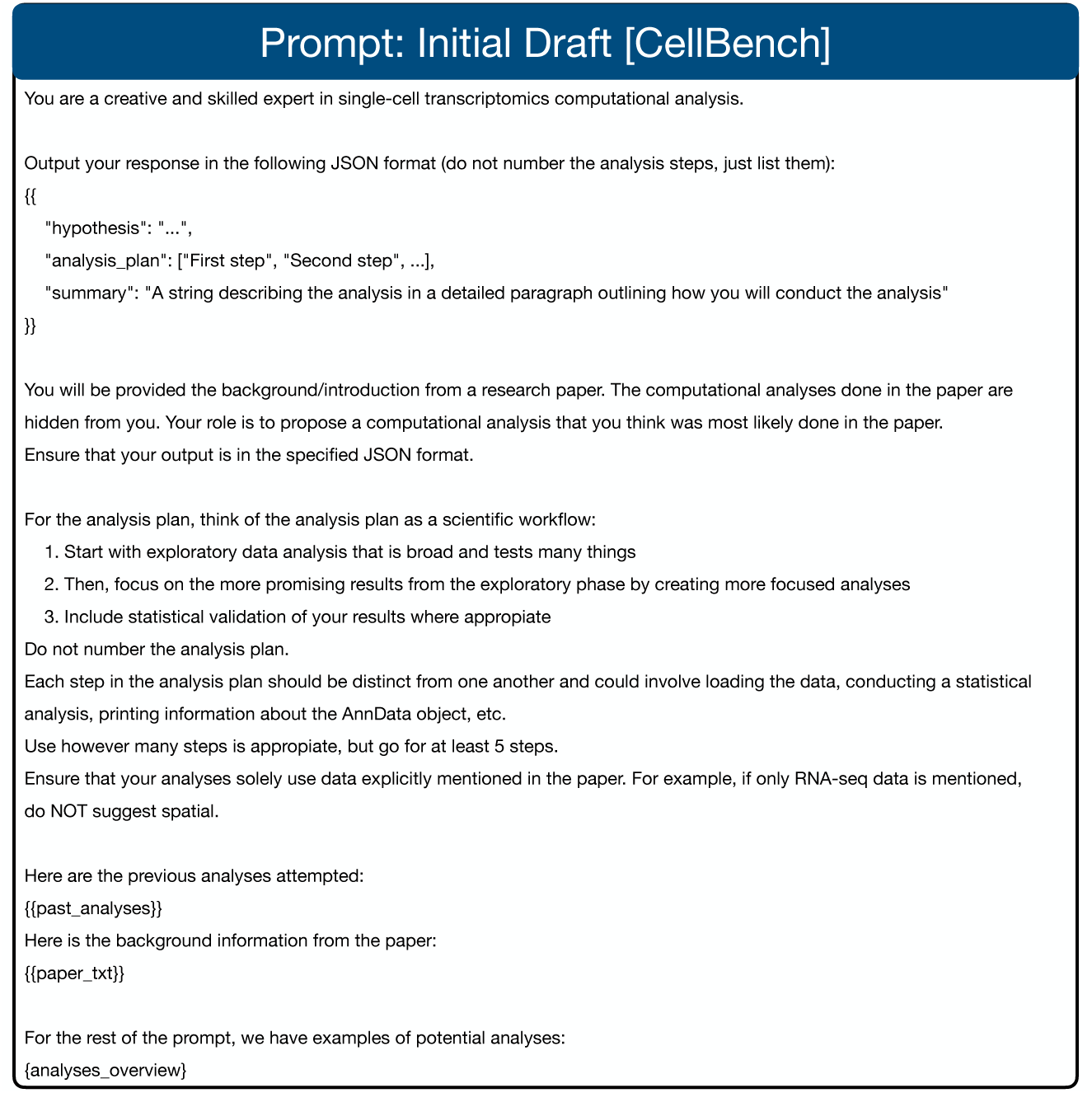
Prompt used for CellVoyager generating an initial analysis idea for CellBench. The system prompt is included at the beginning. Relative to the original first draft prompt, all coding-related guidelines and prompting was removed. Here, “analyses overview” corresponds to a Deep Research derived summary of possible single-cell analyses.

**Appendix Figure 16:**
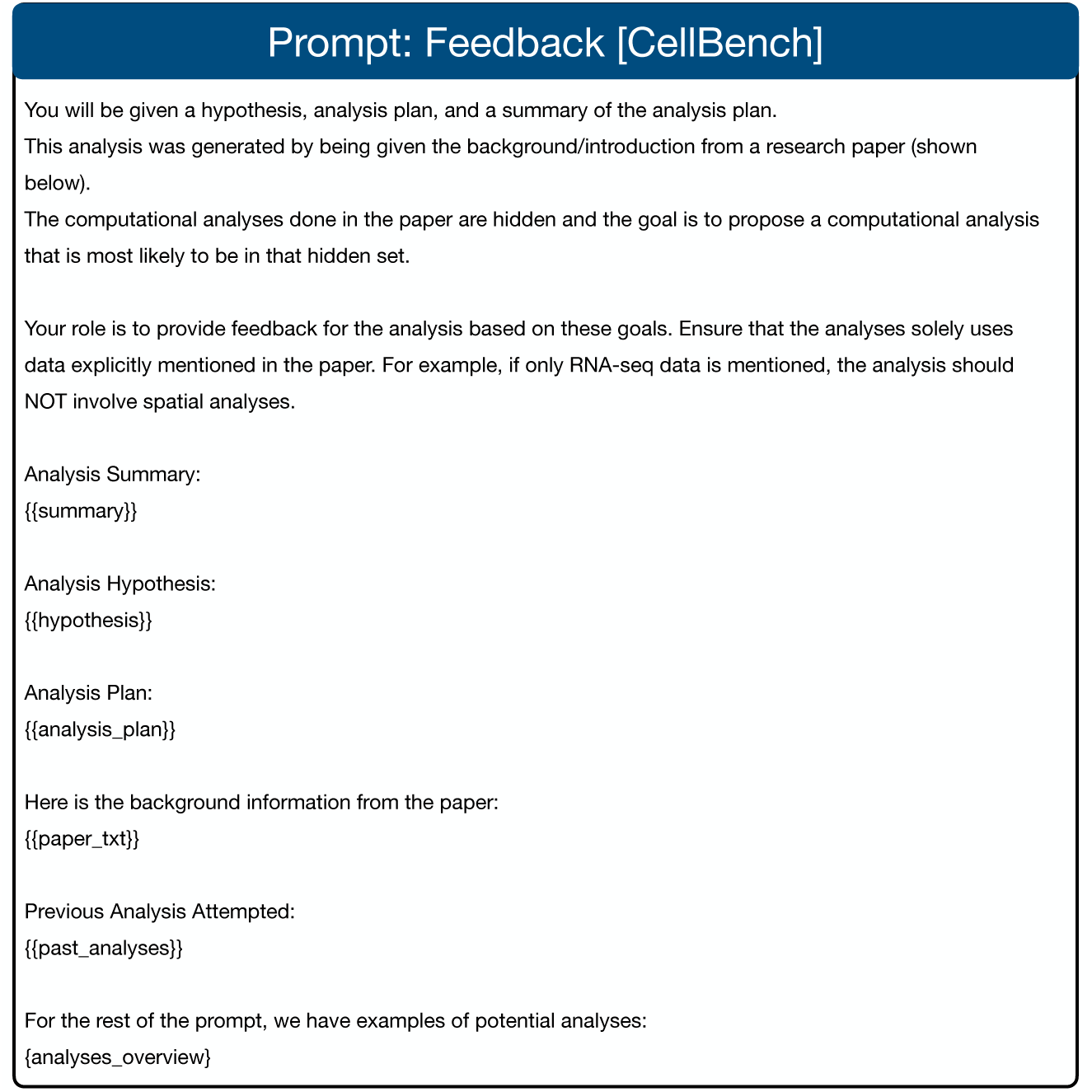
Prompt used for CellVoyager to generate feedback on its analysis idea for CellBench.

**Appendix Figure 17:**
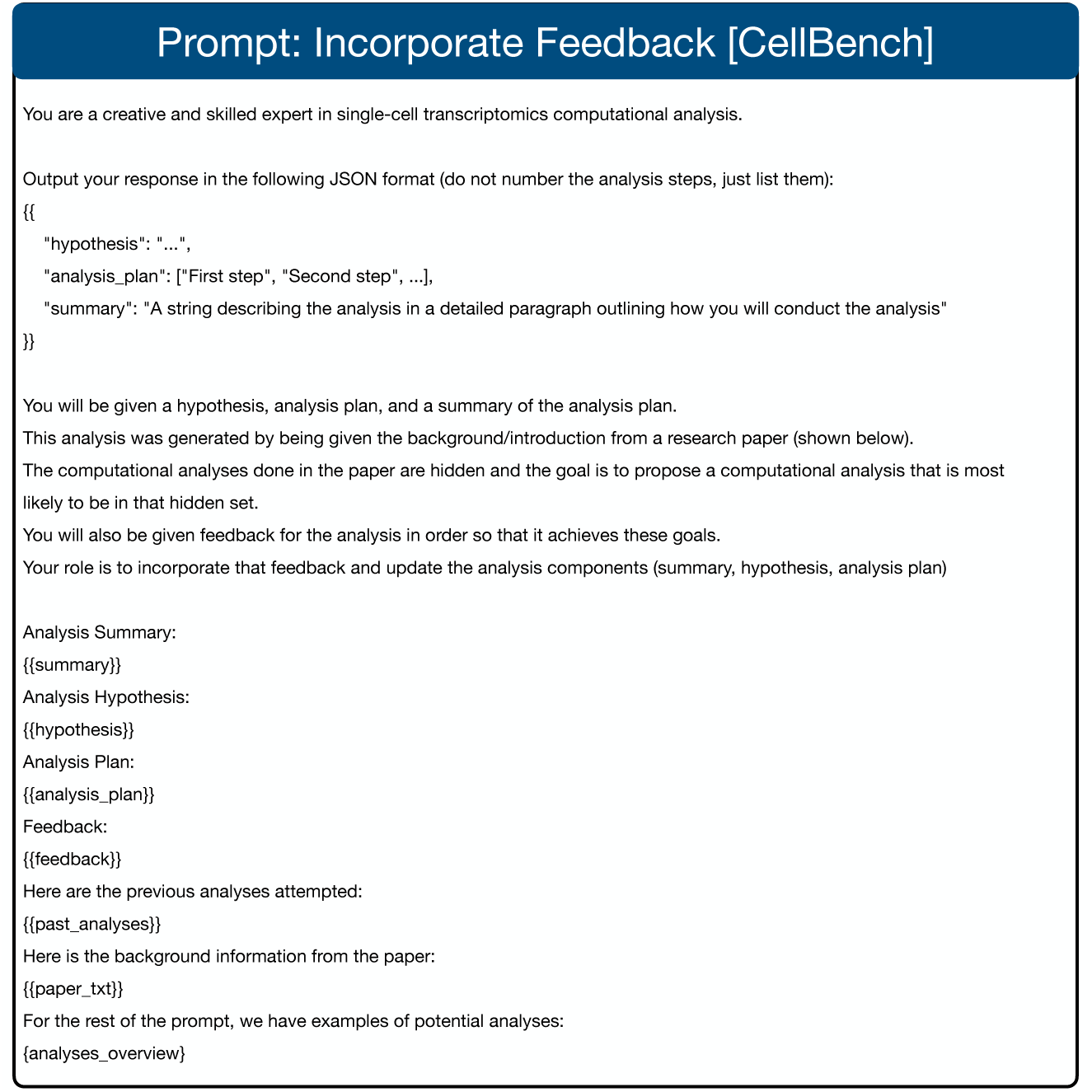
Prompt used for CellVoyager to incorporate its feedback into its analysis idea. Here the system prompt is again included at the beginning.

## B Supplementary Results

**Table 1:** CellBench Results. Micro-averaged and macro-averaged success percentages are shown across 50 papers. Each model was run for 3 times and the mean and standard deviation (in small text) are reported.

| Model | Micro-Avg | Macro-Avg |
| --- | --- | --- |
| GPT-4o | 49.89 (2.73) | 49.41 (2.40) |
| o3-mini | 55.25 (0.80) | 55.46 (0.58) |
| CellVoyager (GPT-4o) | **65.89 (1.81)** | **68.74 (1.52)** |
| CellVoyager (o3-mini) | 64.90 (1.30) | 67.90 (1.47) |

**Appendix Figure 18:**
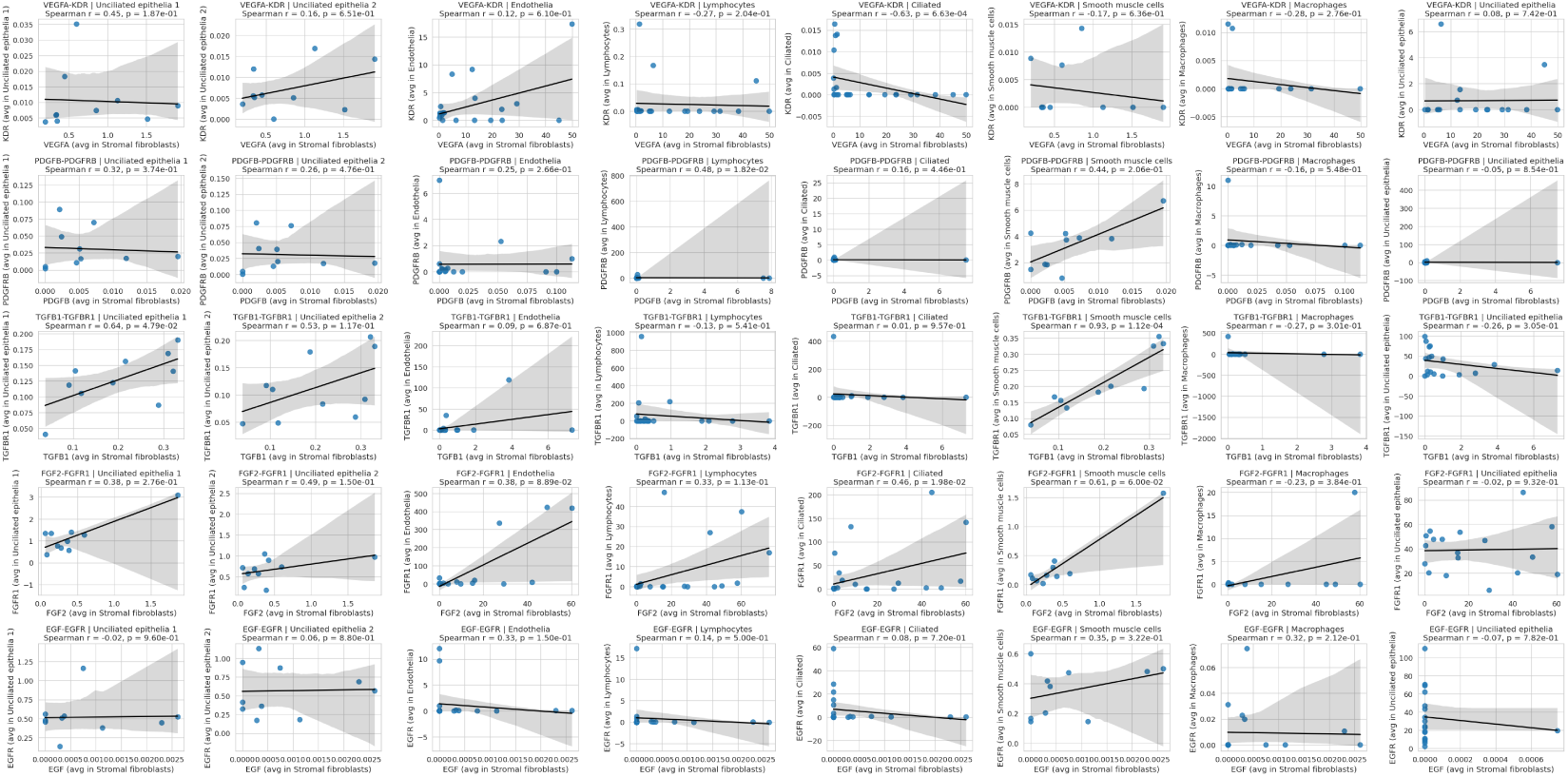
Complete plot from CellVoyager using feedback from the endometrium atlas author.

## Footnotes

1 https://developers.googleblog.com/en/data-science-agent-in-colab-with-gemini/

## Notes

### Competing Interest Statement

The authors have declared no competing interest.

